# Lethal Plague Outbreaks in Lake Baikal Hunter–gatherers 5500 Years Ago

**DOI:** 10.1101/2024.11.13.623490

**Authors:** Ruairidh Macleod, Frederik Seersholm, Bianca de Sanctis, Angela Lieverse, Adrian Timpson, Jesper T. Stenderup, Charleen Gaunitz, Lasse Vinner, Rick Schulting, Olga Ivanovna Goriunova, Vladimir Ivanovich Bazaliiskii, Sergei V. Vasiliev, Erin Jessup, Yucheng Wang, Mark G. Thomas, Russell Corbett-Detig, Astrid K.N. Iversen, Andrzej W. Weber, Martin Sikora, Eske Willerslev

## Abstract

The rise of zoonotic diseases in prehistory is often associated with the Neolithic agricultural transition^1,2^. In particular, plague has been linked to population declines in Late Neolithic Europe^3,4^. Although plague is amongst the most devastating diseases in human history, early strains of *Yersinia pestis,* the causal agent of plague, lack virulence factors required for the bubonic form^5^, and their severity remains unclear. Here, we describe the oldest strains of plague reported so far, associated with two early phases of outbreaks among prehistoric hunter–gatherers in the Lake Baikal region in East Siberia, beginning from ∼5600–5400 years ago (cal. BP). These outbreaks occur across four hunter–gatherer cemeteries; the largest of these (Ust’-Ida I) has a 38.7% detection rate for plague infection (39% detection across all sites). By reconstructing kinship pedigrees, we show that small familial groups are affected, consistent with human-to-human spread of the disease, and the first outbreak occurred within a single generation. Intriguingly, the infections appear to have resulted in acute mortality events, especially among children. Zoonotic transmission is separately indicated by a *Brucella* infection in one of the children. Interestingly, we see differences in functional genomic variants in the prehistoric plague strains, including in the *ypm* superantigen known from *Y. pseudotuberculosis* today. The new strains diverge ancestrally to all known *Y. pestis* diversity and push back the *Y. pestis* divergence from *Y. pseudotuberculosis* by some 2000 years^6^. Our results show that plague outbreaks happen earlier than previously thought and that these early outbreaks were indeed lethal. The findings challenge the common notion that high population densities and lifestyle changes during the Neolithic transition were prerequisites for plague epidemics.

## Introduction

An increased burden of zoonotic diseases in human prehistory has typically been linked to major demographic and behavioural transitions in Western Eurasia, most prominently to the spread of farming and animal husbandry during the Neolithic Revolution^1,2^. The subsistence strategy shift from hunting and gathering to specialised crop cultivation and animal husbandry is associated with intensified resource use and increased sedentism, alongside rapid population growth^7^. This transformation of the human subsistence niche had important consequences for health, associated with changes in nutrition, life-history traits and increased infectious disease burden^8–11^. An increase in zoonosis exposure risk resulting from these lifestyle changes has been interpreted as the cause of a major epidemiological shift in infectious disease burden^1^, with considerable emphasis placed on domesticated animals as novel zoonotic reservoirs^12–14^.

The analysis of ancient pathogen genomes has significantly expanded our understanding of the evolutionary history of human infectious diseases (e.g. *Salmonella enterica*^15^ or hepatitis B^16^), though this has principally been in the context of farming or pastoralist communities. *Yersinia pestis*, the etiological agent of plague, is perhaps the most studied in this regard, and has had devastating consequences upon human populations for millennia. Today, plague is associated with transmission via fleas from rodents which successfully adapted to a human commensal niche in the Neolithic^17^. The detection of early plague cases in Late Neolithic farmers links outbreaks of the disease to a prolonged demographic decline between ∼5300–4900 cal. BP^3,4^, though an alternative explanation attributes the decline to agricultural crisis^18,19^. *Y. pestis* has been estimated to have diverged from *Y. pseudotuberculosis* sometime in the last 50,000 years, based on ancient genomes^4,6,20^.

Studies of prehistoric plague genomes from Late Neolithic and Bronze Age (LNBA) strains predominantly date to between 4700–2400 cal. BP^5,20,21^, and are typically defined as one of two lineages dependent on the presence (LNBA+) or absence (LNBA-) of the *ymt* gene^5^. The presence of this gene determines the survival of the bacterium in the flea digestive tract between rodent and human hosts, and thereby the manifestation of the bubonic form of plague in humans. Lineages of *Y. pestis* that diverged prior to these LNBA clades have also been identified in a Neolithic Swedish individual (5035-4856 cal. BP)^3^, a Latvian individual with Western Hunter–gatherer ancestry (5300–5050 cal. BP)^6^, and in four other Neolithic Swedish individuals (5200-4900 cal. BP)^4^. These genomes lack classic virulence genes (*YpfΦ* prophage and *ymt*), though pangenomic analysis revealed the presence of the locus encoding for *Y. pseudotuberculosis*-derived mitogen (*ypm*), a superantigenic toxin associated with *Y. pseudotuberculosis* (but not later *Y. pestis* strains). This raises intriguing questions over the possible severity of early strains of plague; subsequent LNBA- strains show substantial gene loss, though the virulence potential of these are unknown^5^. Evidence as to the demographic impact of plague infection on prehistoric populations has so far been lacking from these studies.

Middle Holocene hunter–gatherers around Lake Baikal, East Siberia, have been the focus of intensive archaeological study by the Baikal Archaeology Project, yielding important datasets for framing prehistoric hunter–gatherer lifeways^22,23^. These groups demonstrate remarkable continuity of hunter–gatherer lifeways and subsistence, evidenced by an extensive archaeological record of mortuary sites from between c. 8500–3500 cal. BP^24^. The genomes of sampled hunter–gatherers indicate a long-term continuum of Ancient North Eurasian (ANE) and North East Asian (NEA) ancestry until c. 4500–4000 cal. BP^25,26^. By this period, cases of plague from human remains corresponding to the LNBA- strain are documented sporadically among Early Bronze Age burials ^26,27^. Zoonotic spillover events causing plague infections in this region remain a major health concern to this day^28^. These are principally associated with marmots, the primary zoonotic reservoir of plague in this region^29,30^. To explore health and community structure in prehistoric hunter–gatherer groups, we analysed ancient human and pathogen DNA from four cemetery sites in Cis-Baikal (the lake’s western and northern region) dated to c. 5600–5000 cal. BP.

### Outbreaks of Basal Plague Strains

We generated shotgun-sequenced ancient DNA from the archaeological remains of 46 individuals and examined this data for pathogen presence (see Methods). This revealed a conspicuously high occurrence of *Y. pestis* among these individuals, more so than any other pathogen. *Y. pestis* was detected in 18 individuals, indicating two distinct phases of outbreaks of plague infection in four cemeteries. These occur first at Shumilikha (5580–5320 cal. BP) and Ust’-Ida I (5600–5320 cal. BP), and a few centuries later in the sites of Bratskii Kamen (5475–5052 cal. BP) and Serovo (5290–4870 Cal. BP) (see Fig. 1). These sites are all located on banks of the River Angara, a major watercourse draining from Lake Baikal and a vital resource for freshwater fish^31^. At Ust’-Ida I, we also detect the zoonotic pathogen *Brucella*, the cause of brucellosis, in one individual, #26.04. Plague outbreak phases are grouped by the predominant burial-practices at each cemetery: *Isakovo*-style graves in the first outbreak and *Serovo*-style graves in the second phase (see Supplementary Note 1), which are contemporaneous at Lake Baikal between around 6000–5000 cal. BP^31^. This period is defined locally as the Late Neolithic (LN), following Siberian archaeological terminology where the Neolithic is defined on technological criteria such as the introduction of the bow-and-arrow, clay vessels, and stone grinding techniques (i.e. plant and animal domesticates are absent). All four cemeteries were also used during the Early Neolithic (7650–6660 cal. BP) and Early Bronze Age (4970–3470 cal. BP)^31^, though only the LN component of each is considered here.

**Figure 1:**
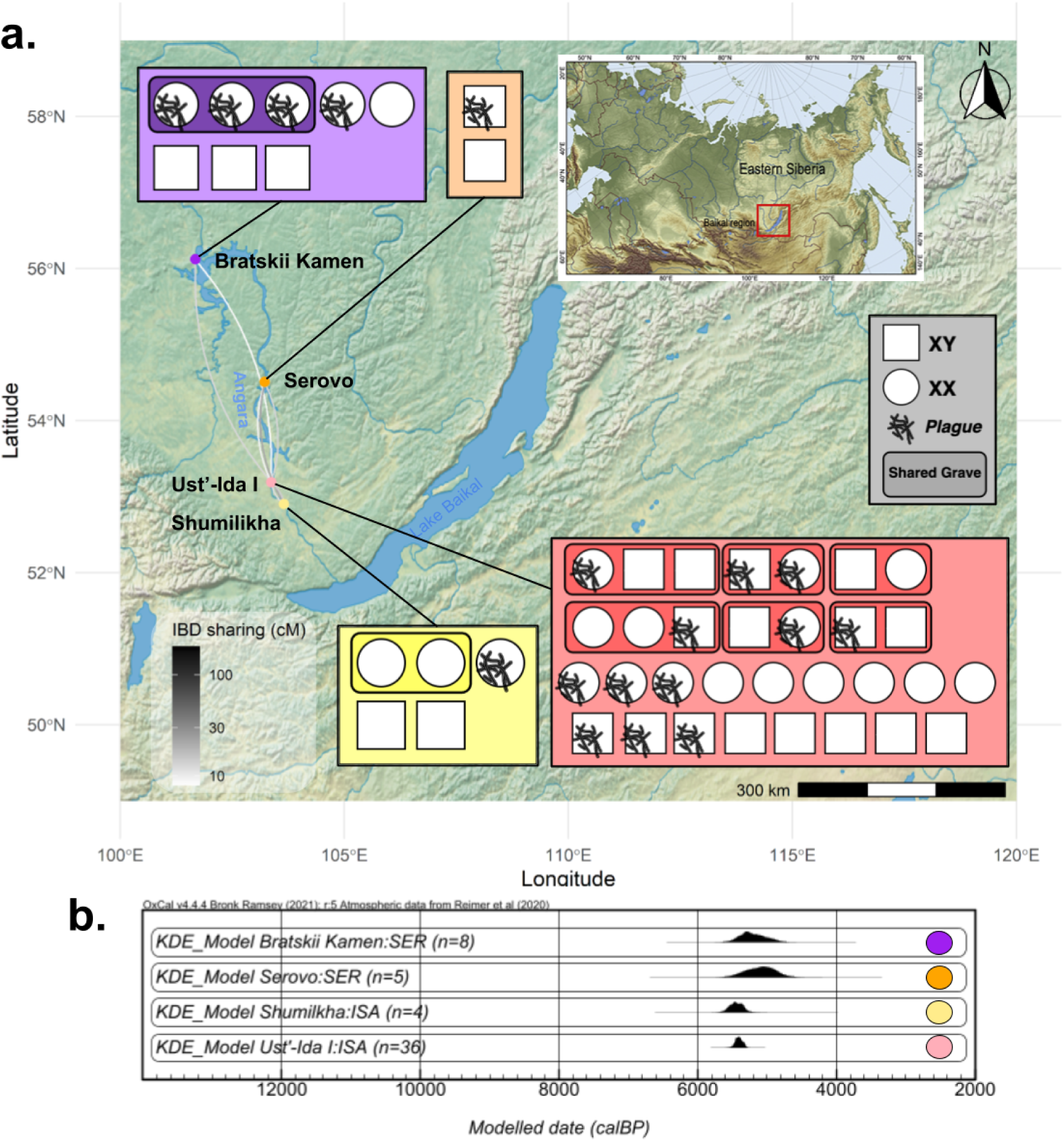
Overview of the spatiotemporal distribution of ancient humans and plague infections in this study. **a**, Locations of affected cemeteries on the Angara River northwest of Lake Baikal, and IBD-sharing between the sampled occupants of cemeteries (pairwise sharing lines between sites, grayscale ramp indicates total IBD-sharing in >3cM segments totalling >10cM in pairwise relationships between sites). Inset are the sampled individuals at each site (31 Ust’-Ida I; 8 Bratskii Kamen; 2 Serovo; and 5 Shumilkha), and those in which plague is detected (**b**, Kernel Density Estimate modelled radiocarbon date distributions for the four cemetery sites (top) and from components of the Ust’-Ida I cemetery specifically (below, after Bronk Ramsey et al.^32^): north and south sector *Isakovo* graves separately; later *Glazkovo* grave groupings (light grey).

Pairwise sharing of Identity-by-Descent (IBD) segments between individuals at these cemeteries indicates recent shared ancestry. Although they are up to 340 km apart, it is entirely probable that mobile hunter–gatherer groups traversed these distances along the Angara River. Very low rates of inbreeding were detected and a high effective population size based on runs of homozygosity was inferred (18,219 individuals, 9,445–42,062 95% CI). This is consistent with the scenario of highly mobile, exogamous hunter–gatherer groups.

Within the hunter–gatherer individuals analysed here, the highest number of detected plague infections was at Ust’-Ida I, which is also the largest *Isakovo* mortuary site in the Baikal region. Here, we found a 39% detection rate (12 out of 31 individuals sequenced), including burials #14 and #56.01, where human genome data was previously reported^25^. Across other sites, we identify one high-coverage plague genome at Shumilikha, four lower coverage genomes from Bratskii Kamen, and one medium coverage genome from Serovo. Overall, we observe a 39% detection rate across LN individuals at these cemeteries (from dental cementum). In comparison, quantitative PCR screening of known Mediaeval plague victims at Smithfield, London^33^ returned a detection rate of 5.7% from bone and 37% from dental pulp tissue (overall 20%), indicating a high rate of false negative plague detection using ancient DNA. To prevent misrepresentation of data, all ancient individuals with screening data from the affected sites are reported here (human autosomal genome coverage ranges from 0.001x to 1.9x, average 0.65x). Direct radiocarbon dates were obtained from nearly all the individuals within the LN components of these cemeteries (a total of 58, including those previously reported from Ust’-Ida I^32^).

*Y. pestis* genomes identified between the two Baikal phases of outbreaks were found to diverge ancestrally to the current known clade of ancient and modern plague strains (Fig. 2). This phylogeny was built using genomes obtained from Shumilikha Burial #34 (6.4x coverage)) from the first phase, and from Bratskii Kamen Burial #22 (1.6x) and Serovo Burial #10 (1.0x) from the second phase. Eight lower coverage genomes were phylogenetically placed using UShER^34^. Through Bayesian inference of node dates in this comprehensive phylogeny of *Y. pestis* and *Y. pseudotuberculosis* diversity, we estimate the origin of *Y. pestis* at a mean of 9317 years ago (95% CI: 7289−11,344)., This pushes back the *Y. pestis* clade diversification date from a previous estimate of 4810–5122 years ago^35^, as would be expected by including *Y. pestis* genomes older than this range. Our result provides a significantly narrower confidence interval than previous estimates based on ancient genome data, which have ranged from 6000 to 50,000 years ago^20^, and revises an earlier estimate of 7400 years ago^6^. The basal *Y. pestis* lineage appears to have arisen as a clone of the ancestor of the O:1c serovar (ENA accession: SAMEA7160327) of *Y. pseudotuberculosis*.

**Figure 2:**
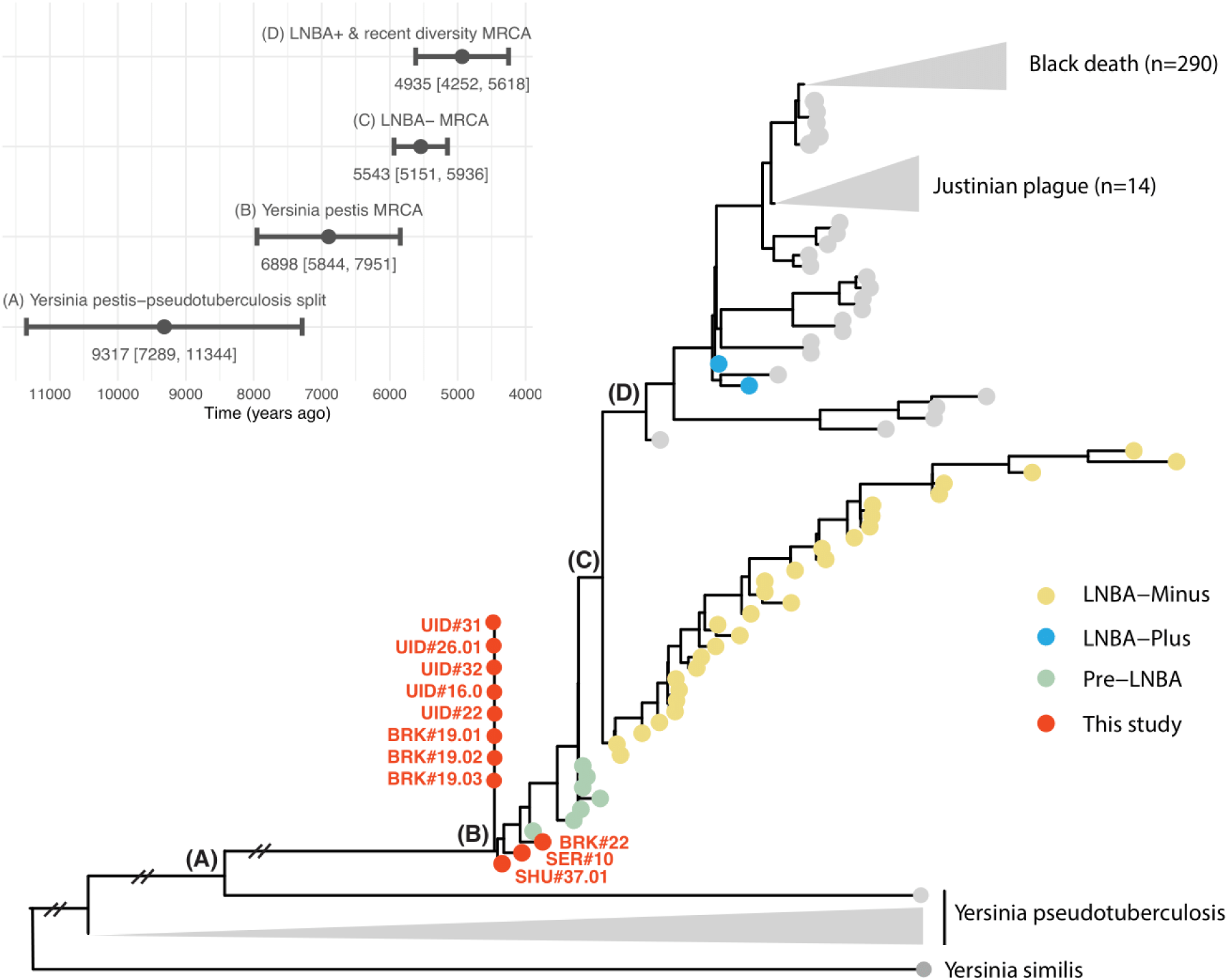
Phylogenetic relationships and molecular dates between hunter–gatherer plague samples from this study and previously published data. The three samples with higher coverage were incorporated directly into the construction of a RAxML phylogeny, whereas the eight lower coverage samples were phylogenetically placed afterwards, all of which shared their most parsimonious placement at the most basal *Yersinia pestis* node. This placement at this node does not constitute a branching position, hence their inclusion adjacent to this node. Divergence times were estimated using BactDating and are shown in the top left panel. BRK = Bratskii Kamen, SER = Serovo, SHU = Shumilikha, and UID = Ust’-Ida I.

Between the two phases, we observe small genetic differences between strains in distinct private mutations in the first and second phase strains (with strict filters for genotype calling, see Methods); this is also clear from the position of nodes in Fig. 2. Although mutation rates in *Y. pestis* are known to be highly variable within different lineages^35^, this result is consistent with a scenario of distinct strains resulting from separate zoonotic spillover events from a local animal reservoir.

### Prehistoric Hunter–gatherer Plague Mortality

To contextualise these plague outbreaks, we considered biological kinship patterns, burial treatment, and age-at-death within the affected hunter–gatherer cemeteries. At the site with the highest positive detection of plague (and largest sample), Ust’-Ida I, radiocarbon dates are exceptionally tightly clustered for a relatively large cemetery^32^, with modelled date ranges indicating that all of the *Isakovo* burials were contemporaneous (Fig. S12). By reconstructing close familial pedigrees, we find that the relationships and ages of family members are consistent with a mortality event over the timespan of less than a single generation (Fig. 3). None of the age-at-death/relationship pairings indicate, for example, children that reached a similar age to their parents, or siblings and half-siblings with very different ages (the greatest sibling age gap is nine years, separated by a middle sibling). Where multiple generations are present, their inferred age-at-death ranges are generally consistent with those expected if all relatives had died at approximately the same time (for example, a 12–15 year old has a 35–50 year old father).

**Figure 3.**
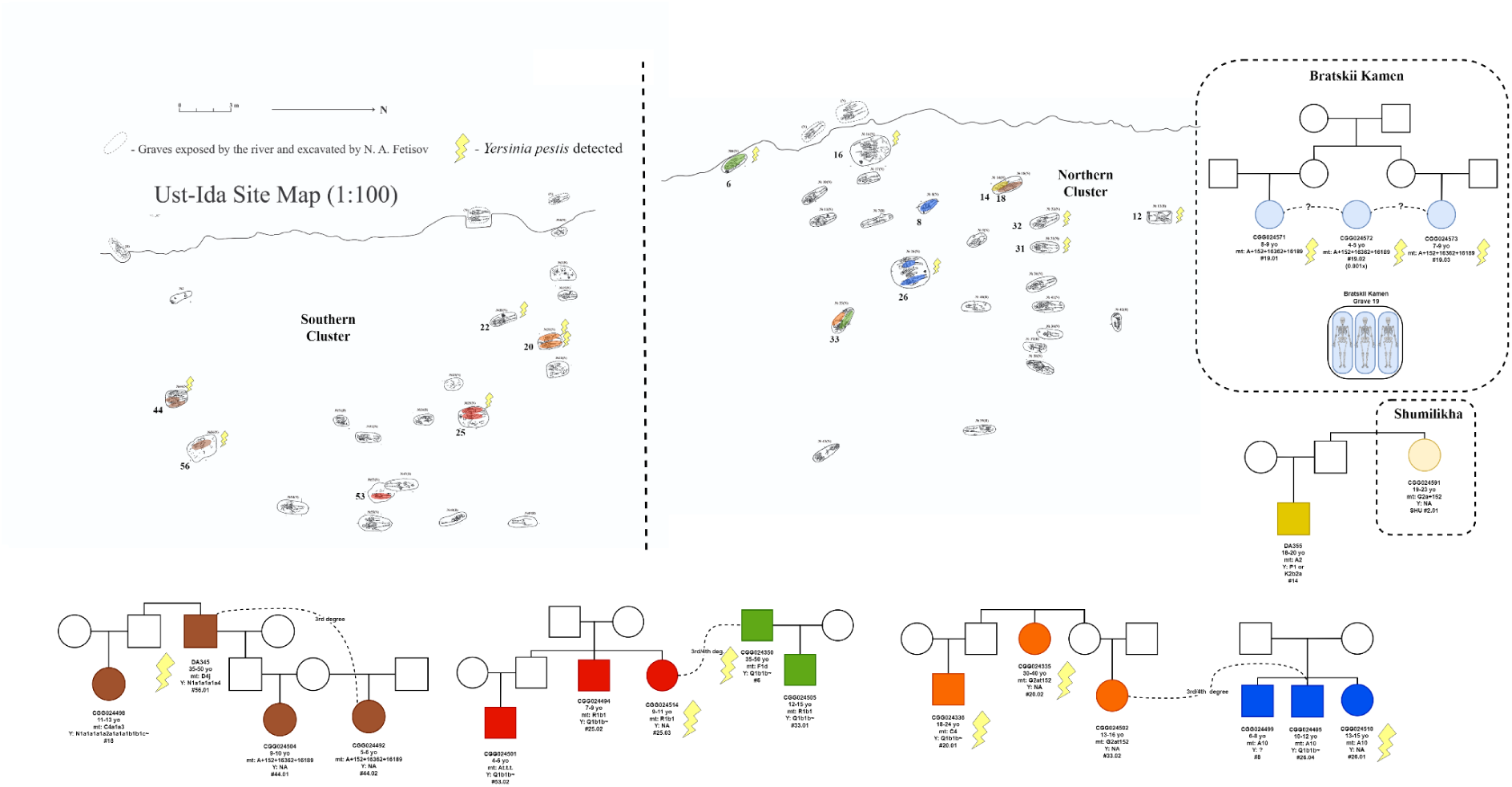
Familial pedigree groups identified from ancient genomes and the site plan for the Ust’-Ida I cemetery. Individuals detected for plague are marked with lightning symbols. Pedigrees are drawn from 30 sampled individuals at Ust’-Ida, five sampled from Shumilikha, and eight sampled from Bratskii Kamen; only close familial relationships are shown; though many other 3rd or 4th degree relationships are detected at Ust’-Ida I (see Supplementary Note 2). The two samples from Serovo, not shown, were found to be 4th degree relatives.

The *Isakovo* mortuary group at Ust’-Ida I is unusual in several other ways among Cis-Baikal hunter–gatherer cemeteries. In addition to the tightly clustered direct radiocarbon dating of burials, childhood mortality is disproportionately high (also observed at Bratskii Kamen, see Fig. 4), and there is a high incidence of multiple interment graves (over half at the site) without any evidence of subsequent grave opening and addition of new burials. This suggests co-occurence of deaths within shared graves; all suggestive of a catastrophic mortality event. Furthermore, an avuncular relationship (yellow pedigree, Fig. 3) exists between the Ust’-Ida I and Shumilkha cemeteries, which are only 37 km apart along the Angara and we can reasonably infer that these groups were in close contact at this time point (matching the concurrent plague outbreak).

**Figure 4:**
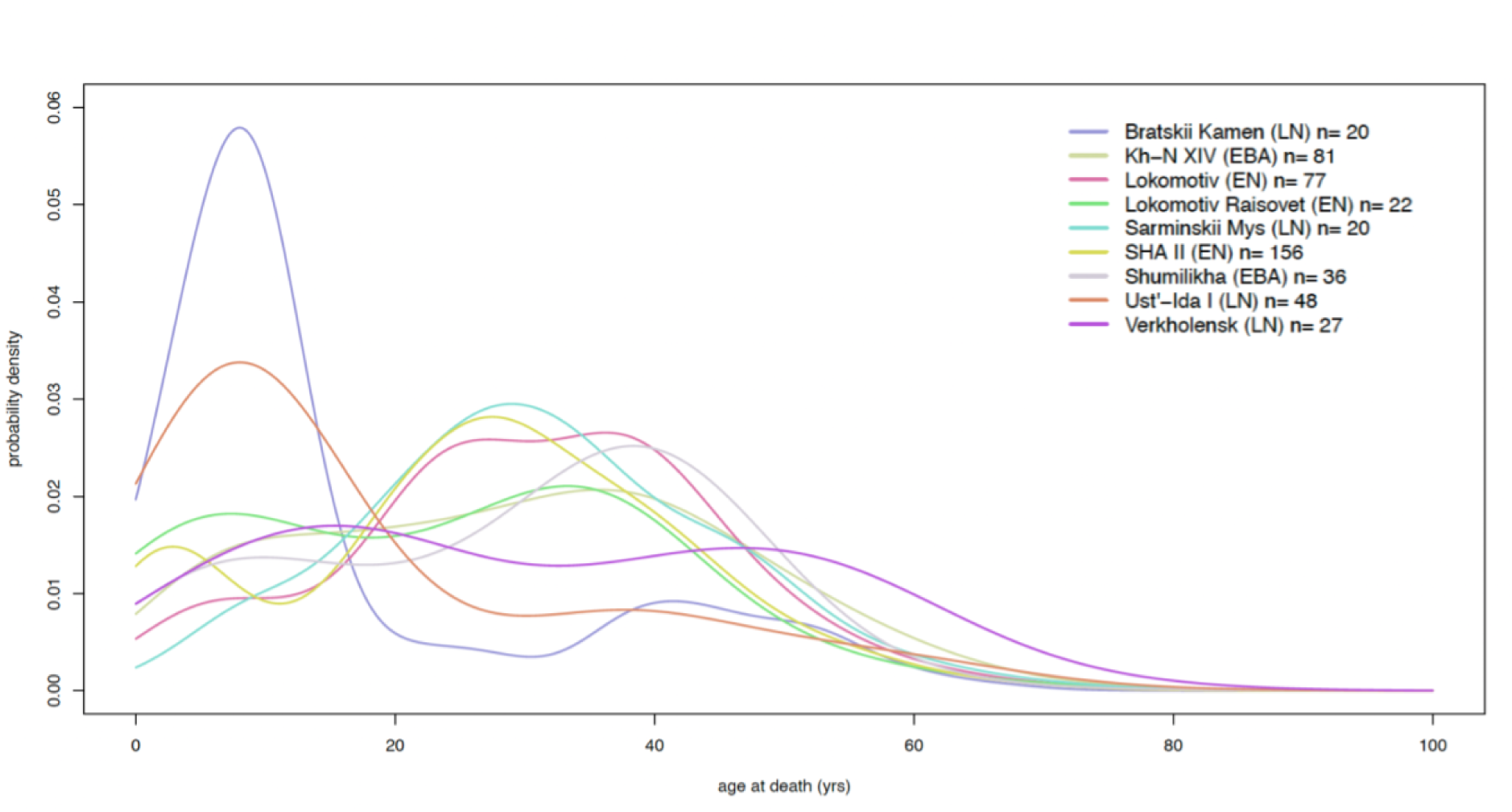
Mortality profiles at Bratskii Kamen and Ust’-Ida I compared with other Baikal hunter–gatherer cemetery populations. Kernel density plot of modelled age-at-death probabilities, based on a null model of a continuous probability of death at any age (see Supplementary Note 5). All relevant assemblages of human skeletal remains from the Cis-Baikal region studied by A. Lieverse with >20 individuals are shown. Sample sizes for sites depicted: Bratskii Kamen LN = 20; Khuzhir-Nuge XIV EBA = 81; Lokomotiv EN = 101; Shamanka II EN = 156; Shumilikha EBA = 35; Ust’-Ida I LN = 48; Verkholensk LN = 27.

In terms of plague detection within grave groups, affected individuals appear conspicuously associated in a number of cases. The burial at Bratskii Kamen features a shared grave of three young girls, aged between 4 and 9 (Fig. 3, left), with similar radiocarbon dates (see Supplementary Note 4). Two of them (#19.01 and #19.03) were inferred as third degree related (most likely cousins); the third had insufficient DNA preservation to confidently infer relatedness, but all three shared a mitochondrial haplotype with three rare private mutations and so were likely close maternal relatives. Genome data for *Y. pestis* were identified in all three, suggesting an outbreak of plague infection in a family, with synchronous deaths of the three children. Similarly, at Ust’-Ida I, a nephew and aunt (#20.01 and #20.02) are buried in a shared grave, with *Y. pestis* identified from both (orange pedigree, Fig. 3). The teenage niece of the aunt, however, is buried in a different shared grave with an unrelated teenage male (possibly suggesting non-biological kinship); his father in turn (green pedigree) is buried in an entirely separate grave, with plague also detected in this individual.

Additionally, some pairs of siblings who are buried together in shared graves show only one individual detected as positive for plague, as is the case with the siblings in grave #25 (red pedigree, Fig 3). In another example, for a sister and brother, #26.01 and #26.04 respectively, the sister is inferred as positive, while the brother is not (though *Y. pestis* reads are detected at just below the threshold for confident identification, Supplementary Data).

These observations are consistent with high false negative detection rate in the palaeogenomic analysis of plague^33^. The brother was also infected with likely non-lethal brucellosis (Supplementary Note 3). In several cases close family members are found in different graves within the cemetery, for example, burial #8, the third sibling of the pair in Grave 26. A pattern is visible where two closely related family members are buried together, while another or others are buried quite far away. This may be consistent with a more drawn out sequence of deaths instead of a single mortality event, if shared graves indicate concurrent deaths, reflecting a scenario of delayed person-to-person disease transmission. No causes of death were apparent other than genetically detected plague infection (though other microbes detected might reflect bacterial coinfections at the time of death, see Supplementary Note 3). Notably, survivors must have existed to bury the deceased, with the typical *Isakovo* mortuary treatment and grave goods, as well as acknowledgement of biological kinship, suggesting a more prolonged sequence of mortality events.

### Epidemiological Implications

A striking aspect of the osteological age-at-death data at the two cemeteries with multiple instances of plague detection is that the demographic profile is highly skewed towards childhood mortality (Ust-Ida I and Bratskii Kamen). Conversely, cemeteries wherein only one affected individual is detected (Shumilikha and Serovo) do not show such profiles (Supplementary Note 1). Ust’-Ida I and Bratskii Kamen both show a peak in mortality at the age range of 7.5–11 years, i.e., in children before puberty (see Supplementary Note 1). In an analysis of mortality profiles across mid-Holocene Cis-Baikal hunter–gatherer cemeteries, these two cemeteries are clearly outliers in terms of the proportion of childhood deaths (Fig. 4). This result was found to be highly statistically significant given a null model of mortality profiles (Supplementary Note 5). Conversely, the 20–25 years age range shows the lowest mortality at Ust’-Ida I, and deaths between 20–35 are completely absent at Bratskii Kamen (Fig. S4). Parents are also conspicuously absent from the pedigree groups; although there are numerous sibling and cousin relationships, there is only one instance of a parent-offspring relationship. The sex ratio in these individuals appears unaffected however (22 XY; 24 XX).

In the context of widespread infection with a plague strain of uncharacterised virulence, this observation could be interpreted as related to differences between child and adult responses to infection. We propose two possible explanations. Firstly, children could be at a greater risk of death due to inherent differences in immune responses between adults and prepubescent children. Secondly, the adults might largely consist of those who had already been exposed to and recovered from the plague as children. Consequently, they would have had some protective immunity against the plague, preventing reinfection or death. The latter would imply that zoonotic infection was a recurring event, which does not match our findings here, that indicate a few short, intense outbreaks with considerable mortality. This explanation would also imply that older individuals would be more likely to have acquired immunity, yet mortality actually increases slightly after the age range of 20–35 years. Therefore, it is plausible that these mortality profiles reflect the age-dependent pathogenicity of this plague strain, although gene functional studies would be needed to substantiate this hypothesis.

At Baikal, the principal zoonotic reservoir of plague is the marmot (*Marmota sibirica*), and marmot hunting for meat and fur has resulted in perennial plague infections especially in young men, exposed during skinning and butchery ^36^. Since the 19th Century, marmots were the most targeted game species by indigenous hunters in this region, originally by trapping^37^, and there are extensive historical accounts of ‘tarbagan plague’ from consumption of infected marmots around Lake Baikal^38^. Prehistoric hunter–gatherer marmot procurement is clearly evidenced by the presence of numerous marmot teeth as grave goods in Early Neolithic *Kitoi* graves^23,39^, though these have not been found in LN graves. Consumption of raw or undercooked marmot organs results in the septicaemic form of infection following the faecal-oral transmission route, while close contact with marmots infected by present-day *Y. pestis* strains causes bubonic or pneumonic infection (or often both), with the latter often occurring secondarily to septicaemic infection^40^ or inhalation of infectious blood droplets during, e.g. skinning^41^.

While a difference in mortality due to behavioural differences between age groups (for example, division of group tasks or roles by age, resulting in higher childhood exposure to marmots) cannot be ruled out, there is little analogous precedent for this with regard to marmots specifically. Furthermore, given the lack of observed heightened childhood mortality in any other Baikal hunter–gatherer cemeteries (Fig. 4), it is unlikely that heightened childhood mortality is the outcome of cultural practices. A more likely explanation for the predominance of childhood mortality (particularly at Ust-Ida I) appears to be age-dependent immune responses that protect children less well than adults against or during infections with gram-negative bacteria, similar to the epidemiological profile of *Y. enterocolitica* and *Y. pseudotuberculosis* infections today^42^. The immune responses in infants and children 10 years of age or younger are immature and characterised by lower levels of bactericidal or permeability-increasing proteins in polymorphic neutrophil granules and B cell immaturity resulting in lower quantities of IgG2 isotypes. These features of the evolving immune system reduce the likelihood that the responses will eliminate infected cells and/or destroy the gram-negative bacteria^43^. This natural increased susceptibility to gram-negative bacteria most likely would include *Y. pestis.* This suggestion is supported by a high incidence of other gram-negative bacteria in childhood pneumonia and sepsis, which decreases with age^44^.

### Functional Variation in Basal Plague Strains

The evolution of *Y. pestis* lineages is shaped considerably by processes of gene loss^45^, a pattern typical of pathogenic bacteria in the transitional process to obligate parasitism, which has also been identified across the LNBA- plague strains^5^. From analysis of the coverage of the classic plague virulence genes, we find that virulence genes absent in published LNBA- and preLNBA strains from Riņņukalns (RV 2039) and Falbygden^4^ are also absent at Baikal (*ymt* and *YpfΦ prophage*), prohibiting the manifestation of bubonic plague (Figure S10).

However, since this virulence gene analysis relies on traditional single-reference mapping, it is restricted to genetic content present in the modern reference. To characterise possible ancestral *Y.pseudotuberculosis* variation in the Cis-Baikal strains which might contribute to our interpretation of their pathogenicity, we mapped sequenced reads to a pan-genome variation graph representing genetic diversity across 82 complete assemblies of the *Y. pseudotuberculosis* species complex (56 *Y. pestis*, 24 *Y. pseudotuberculosis* and one *Y. similis*, based on^4^). We found that the two plague clusters from Lake Baikal carried similar levels of ancestral *Yersinia* diversity only found in *Y. pseudotuberculosis* and *Y. similis* as other preLNBA strains (Figure 5e). For example, we detected the presence of *ypm*, the gene encoding the YPM (*Y. pseudotuberculosis*-derived Mitogen) superantigen known from modern-day *Y. pseudotuberculosis* strains^46^, and recently observed in preLNBA and LNBA- plague strains^4^. Three alleles of this gene exist in modern *Y. pseudotuberculosis*; *ypmA*, *ypmB*, and *ypmC*, with *ypmA* being regarded as the most virulent form of the gene^47^.

**Figure 5:**
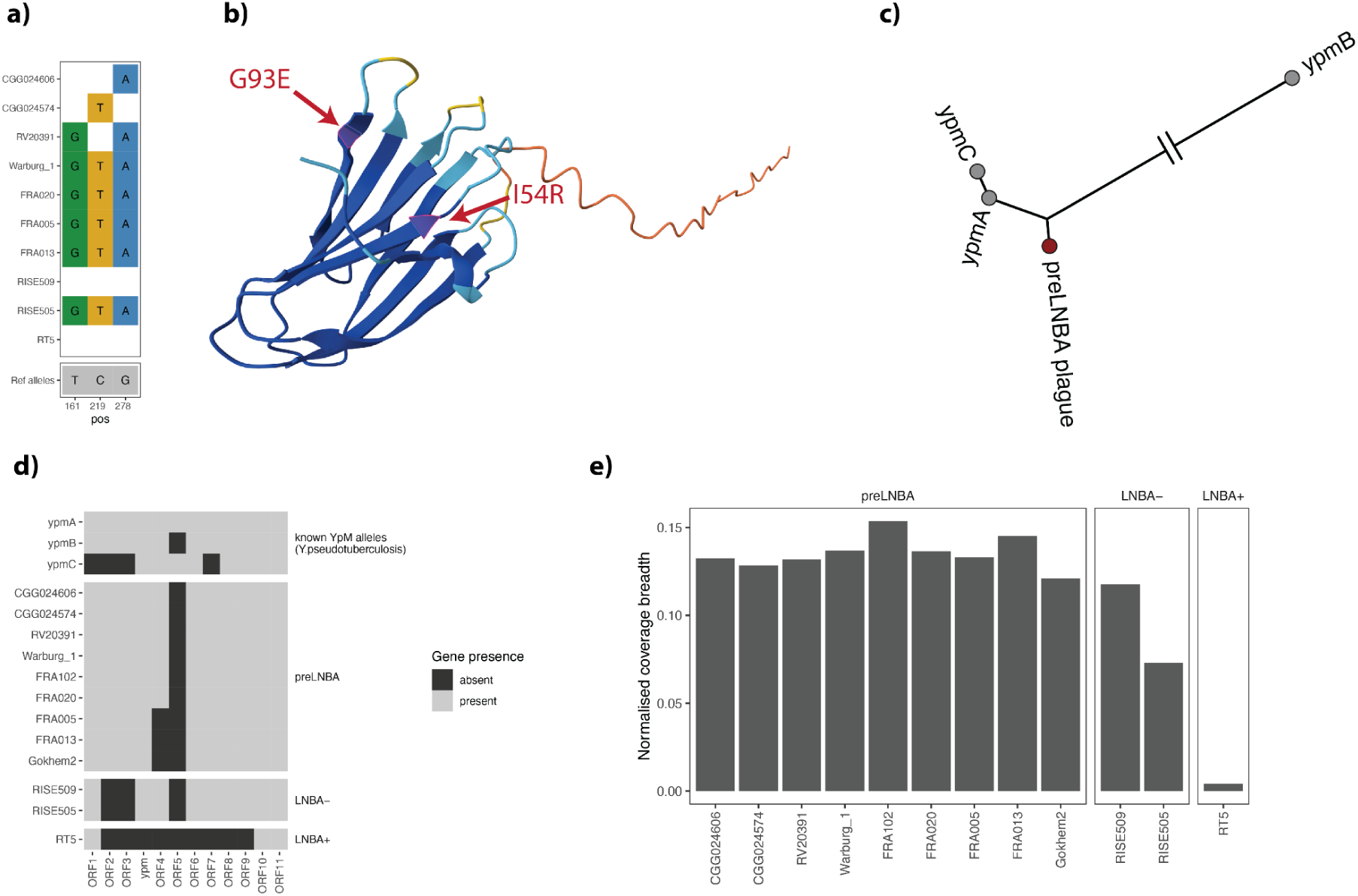
Variation around the *ypm*-locus. **a)** SNPs within the ypm gene. **b)** SNP position (red arrows) on the protein structure of YPMa (PDB accession number 1pm4). This figure was made using data from the AlphaFold database, accession A0A0U1QV71. **c)** Unrooted neighbour-joining tree from the gene sequences of *ypm* variants, including the preLNBA plague form described here. **d)** Open reading frame presence/absence around the ypm-locus. e) Barplot comparing ancestral gene content between prehistoric plague strains from reference graph alignments, using normalised breadth of coverage.

YPM binds to the invariant region of the human leukocyte antigen (HLA) class II molecules and interacts with the variable domain of the β chain of the T cell receptor (TCR). By bridging HLA class II and a TCR, YPM promotes T cell activation and the release of a range of proinflammatory cytokines, further amplifying the immune response^48,49^.

These YPM-associated immune responses have been suggested to be the cause of various inflammatory complications, including encephalopathy, Far East Scarlet-Like fever (FESLF, or Izumi fever in Japan; especially *ypmA*) and a Kawasaki-like syndrome^48,50–52^. Today, FESLF primarily occurs in children younger than 14 and Kawasaki disease mostly in children 5 years or younger. However, Kawasaki disease following *Y. pseudotuberculosis* infection seems to have a higher upper age range. These YPM-related inflammatory complications are likely to also primarily have targeted prepubescent children in the past, further exacerbating the early *Y. pestis* induced morbidity and mortality in the young.

Notably, we find that the *ypm* gene from the two plague strains from Cis-Baikal is closest in sequence similarity to *ypmA* differing at only three base positions: 161 (T>G / isoleucine>arginine), 219 (C>T / synonymous threonine), and 278 (G>A / glycin>glutamate; Figure 5a). These three SNPs appear to be fixed in all plague strains where the gene is present (preLNBA and LNBA- strains; Figure 5a). Since two of these three variants are non-synonymous mutations (I54R and G93E located in distinct beta-sheets of the YPM structure, they could potentially influence protein quaternary structure, protein-protein interactions, and immune response recognition (Figure 5b). Furthermore, by reconstructing the most likely phylogeny of the three known *ypm* variants together with our data, we found that the *ypmB* version of the gene is very divergent from ypmA, C and the gene version from ancient plague. Under the assumption that the root of the tree is located between ypmB and the remaining diversity, the *ypm* from ancient plague appears to diverge ancestrally to both *ypmA* and *ypmC* (Figure 5c).

In addition, we identified 10 open-reading frames (ORFs) around the ypm locus also present in the ancestral form of the plague but absent in later forms. The ORF-region is similar to an unstable region of the *Yersinia pseudotuberculosis* genome with notable low GC content. We found that these ORFs in the Baikal *Yersinia pestis* genomes were similar to those surrounding the *ypmB* variant as reported in other preLNBA strains^4^ (Figure 5d). This pattern with a *ypm* gene similar to *ypmA* combined with a *ypm* locus similar to *ypmB* has, as far as we know, not been observed previously. A possible explanation could be that this diversity in the Lake Baikal preLNBA strains, the closest to MRCA of plague, reflects ongoing local adaptation to marmots and other rodent hosts to a greater extent than humans because the regional animal host reservoir presumably at this time far exceeded that of humans. This, so far unique, combination might affect, e.g., gene methylation and *ypm* transcription levels.

These genetic features of the Baikal preLNBA strains might, together with age-dependent differences in the immune system, partly help explain why prepubescent children predominate among the plague victims although, assessing their true impact requires functional studies

## Discussion

Our findings demonstrate that the earliest known outbreaks of plague occurred in prehistoric hunter–gatherers centuries before infections are observed in Neolithic farmers. These outbreaks were probably the result of zoonotic spillover from wild marmot populations at Lake Baikal. These results support a central/northeast Asian origin for plague, whilst previously the earliest samples had only been reported in northern Europe^4,6^. This is in line with estimates based on the analysis of modern *Y. pestis* diversity^53^. Our phylogenetic analysis reveals that these virulent plague strains are temporally relatively close to the most recent common ancestor of *Y. pestis* and *Y. pseudotuberculosis*, possibly indicating rapid diversification with the transfer to rodent hosts. Additionally, this raises questions around the differentiation of taxa within the *Y. pseudotuberculosis* species complex (which includes *Y. pestis* and *Y. similis*) which ancient genome data alone may not be adequate to answer (given conventional distinctions can be based on pathogenic potential and host range as well). Furthermore, the high lethality of these outbreaks is directly evidenced by mortality profiles and coinciding radiocarbon dates in affected burial sites, indicating that children and adolescents were especially vulnerable; all are insights hitherto lacking for prehistoric plague infections. Until now, the earliest detected strains of plague were of uncertain pathogenicity; their virulence has been the focus of considerable debate, based on genetic data alone^3,5,6,21,54^. Here, we have drawn on multiple lines of evidence from mortuary sites affected by plague (including plague genomes, biological kinship patterns, plague transmission associations, mortality profiles and modelled radiocarbon date ranges) to integratedly characterise the lethal consequences of infection at this period.

The context of these outbreaks is significant to the interpretation of health and epidemiology in the past. That these outbreaks occur in small, mobile prehistoric hunter–gatherer groups emphasises that increased population density, animal domestication and lifestyle changes resulting from the Neolithic transition are not necessary conditions for significant zoonotic outbreaks. This further revises interpretations of plague as a unique contributor to demographic contraction during the Neolithic decline in Europe, as previously suggested^3,4^, especially given the apparent severity of outbreaks identified here. Our findings further reveal insights into the social dimension of these communities during outbreaks, evidencing care for the dead (from the co-interment of close relatives and apparently contemporaneous victims), and the patterns of within-family transmission which evince interpersonal contact in life. This evidence of person-to-person transmission contrasts with previous expectations for basal plague strains^6^.

Significantly, children seem to have borne the brunt of lethality from plague infections at Cis-Baikal. This is possibly the result of differences in prepubertal versus adult immune responses, combined with the effect of distinct genetic differences between preLNBA and later plague strains (e.g. the unique combination of *ypm* and surrounding ORFs). Different mortality rates among age groups have been observed in historical records of plague outbreaks. Similar to our findings, Parish records from the bubonic plague outbreak in London (UK) in 1603 showed a considerably higher child mortality rate (∼5x higher increase in mortality)^55,56^. It is unclear the extent to which cultural or behavioural factors play a role, as servants were also disproportionately affected; however, one distinct difference between the Cis-Baikal outbreaks and mediaeval bubonic plague epidemic is the probable transmission route. While airborne and faecal-oral transmission might have occurred in both, flea bite transmission associated with bubonic plague is unlikely in the Cis-Baikal outbreaks (given the absence of *ymt*). Spread of infectious droplets or aerosols through coughing is documented as the primary transmission mode of pneumonic plague^57^, matching our findings for human-to-human spread inferred from the kinship and archaeological data above. Notably, our results are consistent with previous interpretations that early *Y. pestis* strains could have presented as fatal respiratory pathogens^54^.

Our results indicate that the earliest observed zoonotic spillover was not a one-off event but re-occurred several centuries later with similar lethality, highlighting the prominence of zoonotic infections in prehistoric societies across many different cultural and environmental settings. Additionally, our identification of brucellosis— the first in a prehistoric human context, to our knowledge—suggests evidence of animal-to-human zoonotic transmission in these groups (infection is acquired by direct contact with infected animals^58^, Supplementary Note 3). Recurrent outbreaks of ancestrally diverged plague strains in Cis-Baikal groups between 5500–5000 years cal. BP further suggests a long history of wild rodents as a perennial reservoir for plague spillover. The plague strains genetically closest to those at Cis-Baikal originate from approximately 5000 km west in northern Europe. Given that there is little evidence for external contact with non-hunter–gatherer groups at this time, this supports the hypothesis that a substantial, continent-spanning rodent reservoir of *Y. pestis* might account for frequent isolated spillover events in the subsequent millennia, instead of continuous human-to-human transmission. Moreover, the potential association of prehistoric spillover with human procurement of marmots at Cis-Baikal emphasises the key role which rodent species other than rats likely played in the composition of *Y. pestis* reservoirs. Some 352 reservoir species have been identified from present-day surveillance, a number of which are ecologically long established (e.g. ground squirrels and gerbils)^59^. The scenario of a persistent prehistoric reservoir aligns both with previous findings of rapid repeated infections of diverged plague strains within the same familial lineage from Neolithic Sweden^4^, as well as at Cis-Baikal.

Taken altogether, our findings underscore the universality of zoonotic infection, given the dramatically different life ways of prehistoric hunter–gatherers from European Neolithic farmers. These insights are as relevant for the challenges now faced by the world today as 5500 years ago, with 75% of new human pathogens emerging from animal transmission^60^. Insights into the evolutionary history of pathogens across periods of dramatic demographic and technological change (before the impact of the Neolithic transition in this case) can provide data to contextualise the major challenges humanity is presently facing, such as the climate change-driven disruption of ecological niches around the world ^61^.

## Methods

### Labwork

Ancient DNA was extracted from the dental cementum of molar or premolar teeth from archaeological skeletal remains studied by the Baikal Archaeology Project. Sampling for aDNA was undertaken in dedicated clean lab facilities at the Lundbeck Foundation Centre for GeoGenetics (Copenhagen) and at the Institute of Archaeology, University College London (London). Cementum was isolated specifically from the roots of teeth^see 62^ using a sterilised handheld rotary saw, and pulverised prior to demineralisation and enzymatic digestion. Sampled aliquots were approximately 50–100mg of material. Extraction, purification and library preparation of aDNA for shotgun sequencing followed the approach described in Allentoft et al.^63^, using a double-stranded library protocol following Margaryan et al.^64^ in the first instance, and the ‘Santa Cruz Reaction’ single-stranded library protocol^65^ for samples with low template DNA content. Concentrations for resulting libraries were obtained using an Agilent FragementAnalyzer and pooled at equimolar concentration for sequencing on Illumina NovaSeq 6000 S4 flowcells (100bp paired-end reads) at the GeoGenetics Sequencing Core (Copenhagen).

For libraries where screening sequencing indicated the presence of *Y. pestis* DNA, in-solution capture enrichment was undertaken. Hybridization capture was performed using the Arbor Sciences myBaits kit following Wagner et al.^66^, using the manufacturer’s High Sensitivity protocol, but only with a single round of enrichment. Pooled libraries from the capture reactions were then re-amplified for 16 cycles and sequenced on the same platform as above.

### Preliminary Bioinformatics

Following base-calling of Illumina data using CASAVA (v.1.8.2)^67^, adapter sequences and polyN tails were trimmed from demultiplexed fastq files using AdapterRemoval (v.2.0). Reads were aligned to the human reference genome GRCh38 using bwa aln (v.0.7.18)^68^ (reference genome hg19 was also used for hapRoH analysis, see below). Aligned reads were converted to BAM files, merged across libraries at sample level, sorted, filtered and indexed using Samtools (v.1.21)^69^, then duplicates identified using MarkDuplicates from Picard (v2.18.7), with the following options in place: ‘OPTICAL_DUPLICATE_PIXEL_DISTANCE=12000 REMOVE_DUPLICATES=false TAGGING_POLICY=All VALIDATION_STRINGENCY=LENIENT’. Duplicate reads were then filtered out using Samtools alongside reads with a mapping quality of <30. Summary statistics for sequencing depth and coverage were generated using BEDtools (v2.23.0)^70^ and pysam (https://github.com/pysam-developers/pysam). Estimation of human DNA contamination and damage patterns were performed at a library level, using contamMix^71^, ANGSD (v.0.940)^72^, and mapDamage2.0^73^.

### Human DNA Analysis

Chromosomal sex was inferred based on the ratio of Y and X chromosome aligned reads, following existing confidence intervals^74^. Chromosomal aneuploidies were not detected. Mitochondrial haplogroups were assigned using haplogrep (v.2.4.0)^75^ following annotation of variants using mutserve (v.1.3.0)^76^. Y chromosomal haplogroups were assigned following the approach in^4^.

For exploratory analysis of ancestry through PCA, pseudohaploid genotypes were called by randomly selecting a variant from a pileup generated with Samtools. Samples were then projected into the variation space obtained from using smartpca^77^ to undertake PCA on 2,086,279 SNPs (filtered for transversions only and with minor allele frequency >0.1%) from a reference panel of ancient Eurasian populations^63^. The latter was lifted over from hg19 using hgLiftOver (https://genome.ucsc.edu/cgi-bin/hgLiftOver). All reported samples were included in the PCA by projection.

Diploid genotypes were called using bcftools (v.1.21), and for the analysis of IBD segment sharing, missing diploid genotypes were imputed using GLIMPSE^78^ (for samples with a minimum autosomal genome coverage of 0.1x) following the approach in^63^, and IBD segments called using IBDseq (v.r1206)^79^, followed by genetic clustering by IBD^63^.

Runs of Homozygosity were detected from homozygous-by-descent segments obtained from IBDseq, and from pseudohaploid data subset to the 1240K SNP positions using hapRoH (v.1)^80^. Biological kinship was inferred using KIN^81^, initially running KINgaroo on filtered BAM files targeting the 2,086,279 SNPs described above. Pedigrees were then reconstructed taking into account the resulting log-likelihood estimates for kinship scenarios, uniparental haplotypes, sex and age-at-death (Supplementary Note 2).

### Screening for Pathogen Taxa

Shotgun sequencing data generated from dental cementum was screened for the presence of known human pathogens using the *pathopipe* workflow (https://github.com/martinsikora/pathopipe/) detailed in Sikora et al.^2^. Reads were classified using a fast k-mer approach, *KrakenUniq*^82^ (v.0.5.8), based on a custom database of human pathogens and environmental microbes. For each genus identified in each sample, pairwise alignments using *bowtie2*^83^ (v.2.5.4) were made for all reads classified to that genus against all available species reference genomes for the same genus. Assignments are then made for the presence of pathogen taxa on the basis of the following detection thresholds: unique read count >30, k-mer rank=1, corrected coverage ratio >0.5, average nucleotide identity >0.97, average number of soft clipped bases <8, based on those applied by Seersholm et al.^4^.

### *Yersinia pestis* DNA Analysis

For samples where plague infection was identified through the pathogen screening pipeline, we carried out traditional single reference mapping with Bowtie2^83^ against the plague reference genome (CO92; GCA_000009065.1), with the parameters ‘-D 20 -R 3 -N 1 -L 20 -i S,1,0.50 --end-to-end --no-unal’. Next, duplicate reads and low mapping quality reads (MQ<30) were removed with samtools^69^, followed by calculations of the average depth of coverage in each sample using BEDtools genomecov^70^. We characterised plague cases based on their coverage as either: 1) tentative detections (<0.01X), lower coverage plague cases (0.01-1x) and higher coverage plague genomes (>1x). Individual variants were called based on a minimum depth of four reads at each position.

For the three higher coverage genomes we called genotypes in a sample-wise manner using HaplotypeCaller from GATK^84^, followed by a subsequent step of joint haplotype calling using GenotypeGVCFs on the merged dataset. Next, we removed low-confidence calls using VariantFiltration (GATK) and converted the dataset to multifasta format using bcftools consensus. In doing so, we applied a mask across regions containing the highest proportions of low mapping quality reads, following the approach in Seersholm et al.^4^ to identify such regions. To maintain the coordinates of the reference genome and for consistency with the other aligned samples, fastq data deposited on ENA for reference genomes for *Y. pseudotuberculosis* and *Y. similis* was downloaded and the reads from these realigned to the *Y. pestis* reference genome GCA_000009065.1 and genotypes called and filtered as described above, yielding a multiple sequence alignment for the full *Yersinia* chromosome with 448 sequences on the coordinates of the reference genome. A phylogenetic tree was inferred from the full alignment file including all reference sequences and the three high coverage samples from this study using RAxML-NG^85^ with the GTR + G substitution model and using the *Yersinia similis* reference genome (SAMEA5779183) as an outgroup. We converted the multiple sequence alignment to a haploid VCF using faToVcf (^34^.) with the *Y. pestis* reference genome NC_003143.1.fa as a reference, then built a mutation-annotated tree object from this VCF and the RAxML phylogeny using UShER^34^.

This new UShER phylogeny maintains the original topology but directly assigns substitutions in the VCF to branches on the tree using the Fitch-Sankoff algorithm^86,87^, so that edge lengths are in units of real substitutions. We used matUtils ^88^ to extract a .json file from the mutation-annotated tree protobuf, which can then be uploaded to auspice.us to create an interactive, online, public plague phylogeny annotated with mutations.

For the 8 lower coverage samples, we called a SNP-only vcf using bcftools, filtering for a minimum mapping quality of 30 and bases, a minimum base quality of 30, and a max depth of 1000. We only kept sites which were variable in the reference panel or in which more than one lower coverage sample had a variant called, and further removed all variant sites in the low mapping quality mask described above, yielding a filtered vcf for the lower coverage samples. This low-coverage vcf was used to phylogenetically place the low coverage samples into the mutation-annotated tree using UShER. All 8 lower coverage samples shared a single, maximally parsimonious placement at the root node of the *Y. pestis* clade.

Lastly, we ran BactDating to obtain molecular divergence dates^89^. We used the UShER mutation-annotated phylogeny for this because BactDating requires edges to be in units of substitutions. We included 437 samples which had reliable dates, fixed the topology and root node, and used 1 million iterations and a relaxed gamma model as suggested in^35^. Convergence was confirmed through the trace file as shown in the Supplement. We report median age estimates and 95% confidence intervals for nodes of high interest in Figure 2.

### Variation Graph Analysis

To characterise the full diversity of ancient plague we built a pan-genome variation graph of all known diversity within the *Y. pseudotuberculosis* species complex (Y. pestis, Y. pseudotuberculosis, Y. similis). We used Pangenome Graph Builder (pggb)^90^ on all available *Y. pseudotuberculosis* complex assemblies from NCBI with assembly level characterised as either ‘chromosome’ or ‘complete’. To ensure correct construction of the graph around the plasmids, we built separate graphs for the chromosome and the plasmids and merged these afterwards using vg tools^91^. Next, we indexed the graph and carried out Giraffe^92^ short read mapping to the variation graph of the data from this study and all publicly available ancient shotgun data. Lastly, we identified graph nodes present in the Lake Baikal plague strains but absent in all modern plague assemblies and classified these based on their presence/absence pattern in *Y. pseudotuberculosis* and *Y. similis*. Each node was classified as either ancestral (present in both *Y. pseudotuberculosis* and *Y. similis*) or either *Y. pseudotuberculosis*-derived or *Y. similis*-derived.

### Age-at-Death Estimation

Age-at-death estimation was based on a variety of established anthropological methods. For non-adult individuals (generally <20 years), it was assessed through dental formation and eruption, epiphyseal and long bone diaphyseal measurements, and epiphyseal union, as summarised in Buikstra and Ubelaker^93 pp. 50–52^ and Schaeffer et al.^94^. Adult age estimation focused on skeletal morphological changes, namely those of the pubic symphysis^95,96^ and iliac auricular surface^97–99^, but also palatine and ectocranial suture closure^100–102^. For all individuals, as many methods as possible were considered based on the state of skeletal and/or dental preservation.

## Acknowledgements

This study was supported by a SSHRC doctoral studentship grant (G101449: ‘Individual Life Histories in Long-Term Cultural Change’ for R.M.) and the Lundbeck Foundation (no. R322-2019-2610 to F.V.S.), and Riksbankens Jubileumsfond (no. M 21-0018). Research reported in this paper has been part of the Baikal Archaeology Project and Baikal–Hokkaido Archaeology Project supported by grants from the Social Sciences and Humanities Research Council of Canada: Major Collaborative Research Initiative Nos. 410-2000-1000, 412-2005-1004, and 412-2011-1001; and Partnership Grant No. 895-2018-1004 as well as numerous matching funding provided by the University of Alberta and other partner organisations. Additional funding was provided by the Government of the Russian Federation, grant No. 075-15-2019-866 (“Baikal Siberia in the Stone Age: At the crossroads of the worlds”). Furthermore, the research at the University of Copenhagen was carried out under the Lundbeck Foundation GeoGenetics Centre, which is supported by the Lundbeck Foundation (nos. R302-2018-1799 2155 and R155-2013-16338), the Novo Nordisk Foundation (no. NNF18SA0035006 and NNF24SA0092560), the Wellcome Trust (no. UNS69906), the Carlsberg Foundation (no. CF18-0024), the Danish National Research Foundation (no. 44113220 and DNRF174) and the University of Copenhagen (KU2016 programme). M.G.T. is supported by ERC Horizon 2020 research and innovation programme grant agreements: no. 951385 (COREX) awarded to M.G.T., no. 865515 (SUSTAIN) awarded to Maria Ivanova-Bieg, no. 324202 (NeoMilk) awarded to Richard Evershed, no. 788616 (YMPACT) awarded to Volker Heyd, and by Wellcome Senior Research Fellowship Grant 100719/Z/12/Z awarded to M.G.T. Data analysis was supported by the NSFC BSCTPES Project (No. 41988101), the CAS Youth Interdisciplinary Team Fund, the National Key Scientific and Technological Infrastructure project “Earth System Numerical Simulation Facility” (EarthLab, 2023-EL-ZD-000111) and nd SMU’s Center for Research Computing.

The authors express their gratitude to Emily Tilby for assistance in drafting figures; Maria Madrona, Marcus Hjorth, Andreas Poersksen and Lærke Kjærsgaard Hansen for assistance in labwork; to Andrea Hiob for management of the Baikal Archaeology Project; to Anna Razeto and Line Olsen for management of the Lundbeck Foundation GeoGenetics Centre projects; to Stephen Shennan and Richard Durbin for useful discussions; and to N. N. Mamonova for providing additional data for age-at-death estimations.

## Contributions

Ancient DNA labwork was undertaken by R.M., F.V.S., J.T.S., C.G., and L.V.

Computational analysis was undertaken by R.M., F.V.S., B.D.D-S., and M.S., and supervised by M.S. (with input on phylogenetic analysis specifically from R.C.-D.).

Archaeological research contributing to this study was undertaken by A.W.W., A.L., R.S., E.J., O.I.G., V.I.B., and directed by A.W.W.

Osteological analysis specifically was undertaken by A.L.

Radiocarbon dating was undertaken by A.W.W. and R.S.

Samples were curated by E.J., S.V.V, O.I.G and V.I.B.

Specific areas of expertise for interpretation and further analysis were provided by A.K.N.I (immunology and pathology); A.T. and M.G.T. (modelling mortality profiles); Y.W. (ecology). The initial draft writing was led by R.M., together with F.V.S and A.K.N.I., with subsequent contributions from B.D.D-S., A.L., E.J., R.S., M.G.T., A.W.W., M.S., and E.W. All authors reviewed, commented on and approved the final version of the manuscript for submission.

This project was undertaken in the context of the PhD research of R.M., which was conceived by A.W.W. and E.W.

Research design for plague analysis was conceived by M.S. and E.W., with input from R.M. and F.V.S. E.W. was primarily responsible for supervising this research.

## Notes

### Competing Interest Statement

The authors have declared no competing interest.

