## Supplementary information for "Lethal Plague Outbreaks in Lake Baikal Hunter–gatherers 5500 Years Ago"

**Supplementary Data Tables**
Supplementary Table 1: *Genetic and archaeological metadata of ancient individuals, and reported radiocarbon dates.*
Supplementary Table 2: *Plague detection results from screening of shotgun sequencing data.*
Supplementary Table 3: *Other pathogen detection results from screening of shotgun sequencing data.*
Supplementary Table 4: *Results of pairwise IBD-sharing.*

#### Supplementary Note 1: Archaeological Context

Hunter–gatherer cemeteries in the Cis-Baikal Region (the area on the northwest side of Lake Baikal including the Angara River Valley and Upper Lena River) have been the focus of intensive archaeological research by the Baikal Archaeology Project since the mid-1990s, though most were first excavated in the 19th and 20th centuries, in particular under the Soviet archaeologist Alexei Pavlovich Okladnikov. Okladnikov defined the distinct cultural attributes of graves typologically as mortuary traditions [1]. His model was later revised based on direct radiocarbon dating of hundreds of human burials by the Baikal Archaeology Project[2,3], resulting in the following sequence: Late Mesolithic (*Khin* mortuary group), Early Neolithic (*Kitoi* mortuary tradition), Late Neolithic (both *Isakovo* and *Serovo* mortuary traditions), and Early Bronze Age (*Glazkovo* mortuary tradition). As noted in the main text, the Neolithic in Siberian archaeology is defined following the Russian archaeological tradition according to which it is defined on technological grounds, that is by the introduction of the bows-and-arrows, clay vessels, and stone grinding techniques. There is no evidence for formal cemeteries from the Middle Neolithic and so it is effectively lacking from this sequence of mortuary sites. The focus of the present study is on the following Late Neolithic (5600–5000 years cal. BP) burial sites: Ust’-Ida I and Shumilikha (*Isakovo* mortuary tradition), and Bratskii Kamen and Serovo (both representing the *Serovo* mortuary tradition). On the basis of previously published stable isotope data, some evidence of dietary trends in these groups have been identified, with the *Isakovo* isotopic sample primarily from Ust’-Ida I (Angara river valley). Stable isotope data for *Isakovo* individuals from Ust’-Ida demonstrate reliance on local middle Angara River fish and terrestrial game[4]. Data for the direct comparison of dietary isotopic trends between Late Neolithic cemeteries in the Angara River valley are limited but some new results have been recently obtained (including BRK, SER, and SHU cemeteries) and will be published in near future.

Okladnikov originally believed the *Isakovo* group preceded the *Serovo*[1], though these are now known to be contemporaneous[5]. *Isakovo* burials are predominantly found along the Angara River (which flows north from Lake Baikal and forms a tributary of the Yenisei), and many burials are oriented parallel to the river with heads pointing upstream although variation in this regard does exist. Bodies are positioned extended and supine, and cemeteries are typically of a small to medium size (fewer than 25 individuals). Most graves comprise single inhumations, though shared graves are not unusual, and may include children in particular. Graves display stone structures and grave goods assemblage commonly consists of clay vessels (mitre-shaped), in addition to lithic arrowheads, bone and antler points or composite tools (fishing gear is rare, despite the proximity to the river, and red ochre is absent).

*Serovo* burials are distributed also along the Angara River but also along the Upper Lena River (in the latter sometimes referred to as the Archaic mortuary tradition[6], as well as the Little Sea microregion. The *Serovo* mortuary tradition is similar to *Isakovo* in terms of cemetery size (with the exception of Ust'-Ida where its Isakovo component is largest known of either tradition anywhere in Cis-Baikal), use of stone structures and extended supine body position, but many burials are instead oriented perpendicular to the river, with the head pointing away (but variations from this pattern are not uncommon), and there are differences in grave goods assemblages. In particular, large bifacially formed lithic spearheads and egg-shaped clay pots appear to distinguish *Serovo* burials from *Isakovo*. Use of red ochre is rare and limited mostly to isolated stains, and there is possibly a higher incidence of rich grave goods associated with *Serovo* burials than *Isakovo*.

Ust'-Ida I

Ust'-Ida I is a large cemetery located on the east bank of the Angara River approximately 250 km north of Lake Baikal, first identified from the discovery of a single grave by A.P. Okladnikov in the 1950s, and subsequently several more in the 1980s. The site was systematically excavated under the direction of Vladimir I. Bazaliiskii from 1987 to 1995, resulting in the discovery of multiple phases of use: Early Neolithic *Kitoi* with 1 grave; 32 Late Neolithic *Isakovo* graves (comprising 48 individuals, the most substantial component of the cemetery), and 18 Early Bronze Age *Glazkovo* graves (comprising 19 individuals). Unfortunately, this cemetery has not yet been published as a monograph and is known only from a short site report[7]. Comprehensive radiocarbon dating of the *Isakovo* component of this cemetery was undertaken by the Baikal Archaeology Project (Weber et al. 2006, Radiocarbon paper; Weber et al. 2021; Ramsey et al.[8], resulting in direct dates for 36 individuals (see Supplementary Note 4). In addition to stable isotope data indicating a dependency on local fishes from the middle Angara[4], activity-induced dental modifications are observed exclusively in males at Ust'-Ida I[9]. These occlusal grooves are linked to the processing of fibres for the production of thread (possibly for the construction of nets). Grave goods are highly repetitive and include clay pots with net impressions, lithic arrowheads, ground adzes and knives made of slate and nephrite, bifacial triangular knives, composite insert tools, bone points of various forms, harpoons, and red deer canine pendants and a rare anthropomorphic figurine[7].

Genetic data have previously been published from this site in an early study of mitochondrial DNA haplotype variability, amplifying a 440bp region of the mitochondrial genome in 19 ancient individuals[10], and subsequently from the mtDNA coding region in 39 ancient individuals [11]. Ancient genome data were previously reported for four individuals [12], corresponding to Burials 22, 30, 56.01, and 14 (published sample numbers DA342, DA344, DA345, and DA355 respectively).

Shumilikha

Shumilikha, also sometimes referred to in the archaeological literature as Ust'-Belaia II (distinct from Ust'-Belaia), is located at the confluence of the Angara and the Belaia Rivers

(Fig. S1). This site was excavated in 1972–1973, and published in a monograph edited by V. V. Svinin in 1981[13]. It predominantly comprises Early Bronze Age graves of *Glazkovo* graves (39 inhumations within 37 graves, 2 instances of shared graves). Additionally, there are 2 single Early Neolithic *Kitoi* graves, and 10 burials across 6 Late Neolithic graves, three of which have been classified as *Isakovo,* although this designation is somewhat inconclusive.

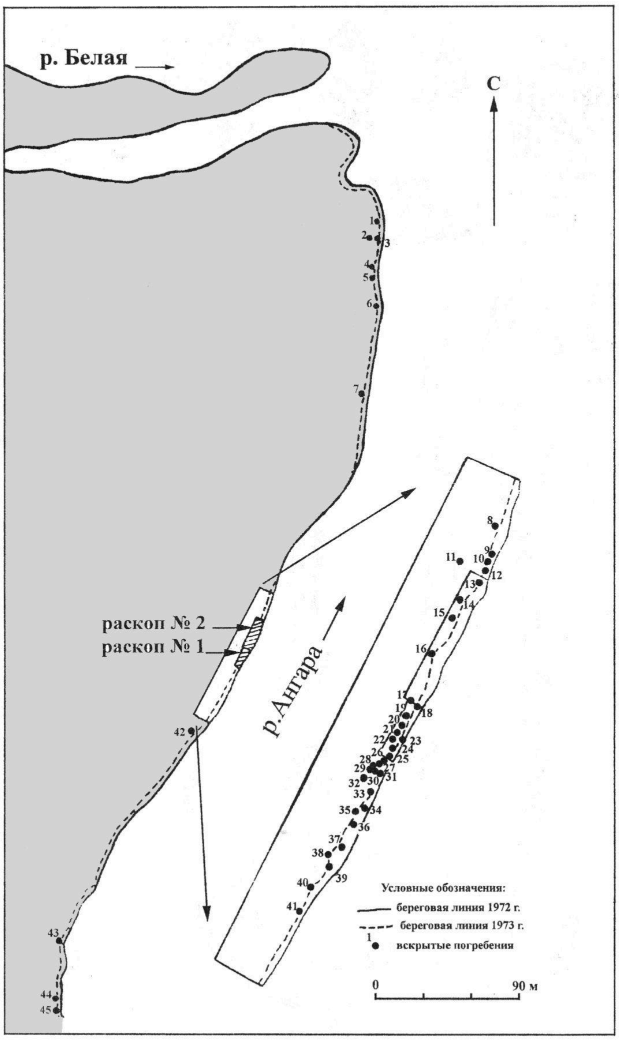


**Figure S1: Site map of excavated graves at Shumilkha, indicating its location on the Angara River.** From Svinin 1981[13], page 53.

<u>Bratskii Kamen</u>

The site of Bratskii Kamen is located south of the city of Bratsk in the now flooded area of the Bratsk Reservoir, which was dammed for a hydroelectric power plant completed in 1967. The site predominantly comprises Late Neolithic graves (21 individuals within 16 graves: 3 *Isakovo* and 12 *Serovo*, see Fig. S2), though also includes several other graves: 1 Early Neolithic, 1 Early Bronze Age *Glazkovo* grave, and 4 additional graves with absent diagnostic characteristics (all single burial graves). The site was initially excavated in 1932, and excavated again in 1955–1956 by A. P. Okladnikov prior to the flooding of the local area [14,15].

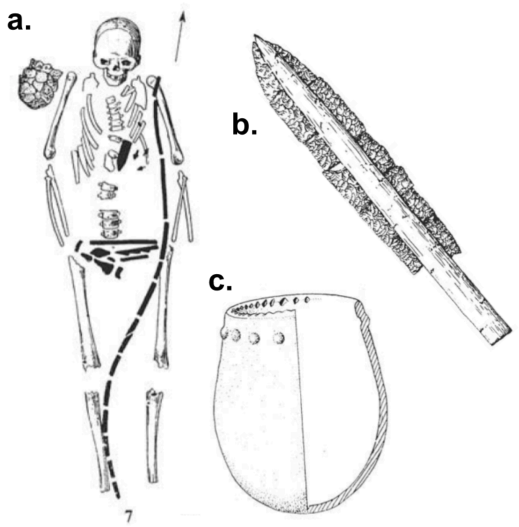


**Figure S2: Features of the *Serovo* mortuary grouping at Bratskii Kamen. a**, plan of Grave 7, featuring a bow above the skeleton; **b**, composite point with lithic inserts; **c**, clay vessel. Redrawn by V. Bazaliiskii after Okladnikov[15].

Serovo

Serovo (sometimes ‘Zverevo’) is the type site upon which *Serovo* mortuary grouping is based, and is located on the west bank of the Angara river, 145 km north (downstream) of Ust’-Ida I, and over 200km south by southeast from Bratskii Kamen. Inhumations at Serovo were first excavated in the 1930s, and the site was extensively excavated in 1957, led by A. P. Okladnikov [15]. Burials are predominantly Late Neolithic *Serovo* graves (16 individuals in 15 graves), though with two Early Neolithic and two Early Bronze Age *Glazkovo* outlier graves, and five missing contextual identification.

Bioarchaeology of Human Remains

Demographic profiles of the Late Neolithic remains from Ust’-Ida I, Bratskii Kamen, and Serovo (Figures S3–S5) reveal the uniqueness of the sites, all being dominated by non-adult individuals (those under the age of 20 years at death). In particular, 65–75% of individuals interred at Ust’-Ida I and Bratskii Kamen represent prepubescent children under the age of 15 years. While Serovo (Figure S5) has relatively fewer non-adult individuals (44%), their proportion is still considerably higher than is typical for the Middle Holocene Cis-Baikal. As a comparison, non-adult individuals from EBA cemeteries in the region, such as Khuzhir-Nuge XIV and the EBA component of Ust’-Ida I, represent 26–33% of those interred[16–18]. Demographic data are not yet available for Shumilikha. Explaining the apparently high mortality of infants and children in the Late Neolithic is not possible from macroscopic analyses of human remains. Unfortunately, the vast majority of pathological conditions—especially acute infections causing death—do not affect skeletal and dental tissues[19]. Indeed, paleopathological data from these four sites reveal low frequencies of skeletal lesions, most representing degenerative changes, such as osteoarthritis, on adult individuals. Lesions consistent with physiological stress or “poor health” reflect chronic infections (e.g., sinusitis) or metabolic conditions, being documented on fewer than 20% of individuals at any site. Paleopathological data are similar among the four sites, and typical, again, for the Middle Holocene Cis-Baikal[16–18].

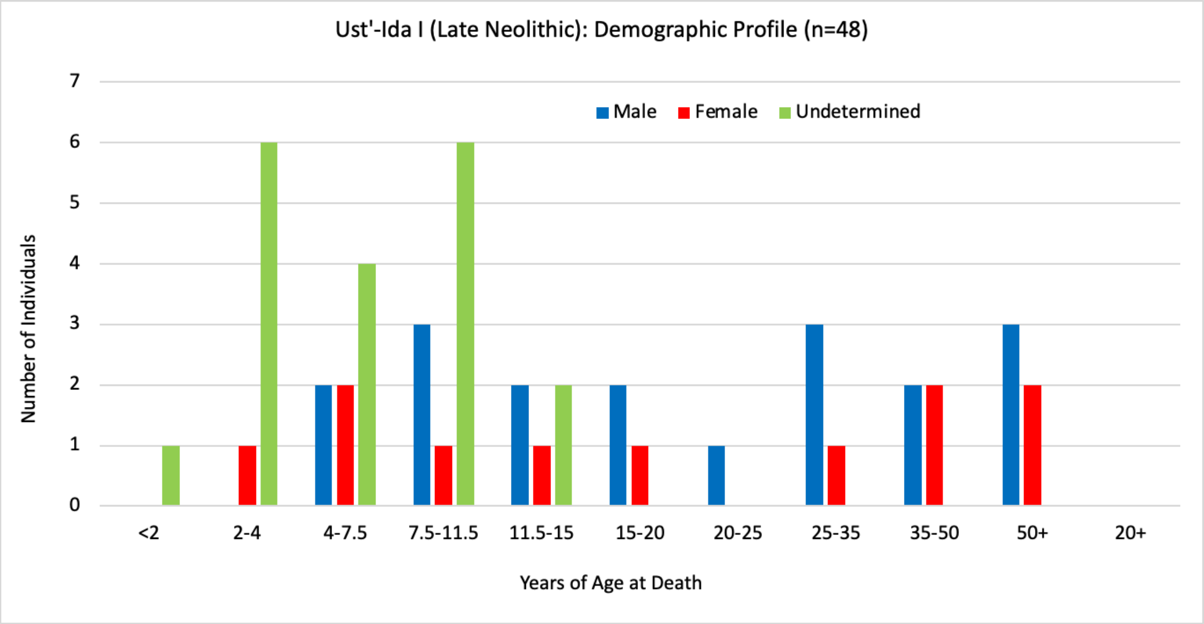


**Figure S3: Mortality profile inferred from age-at-death estimates for Late Neolithic individuals at Ust'-Ida I.**

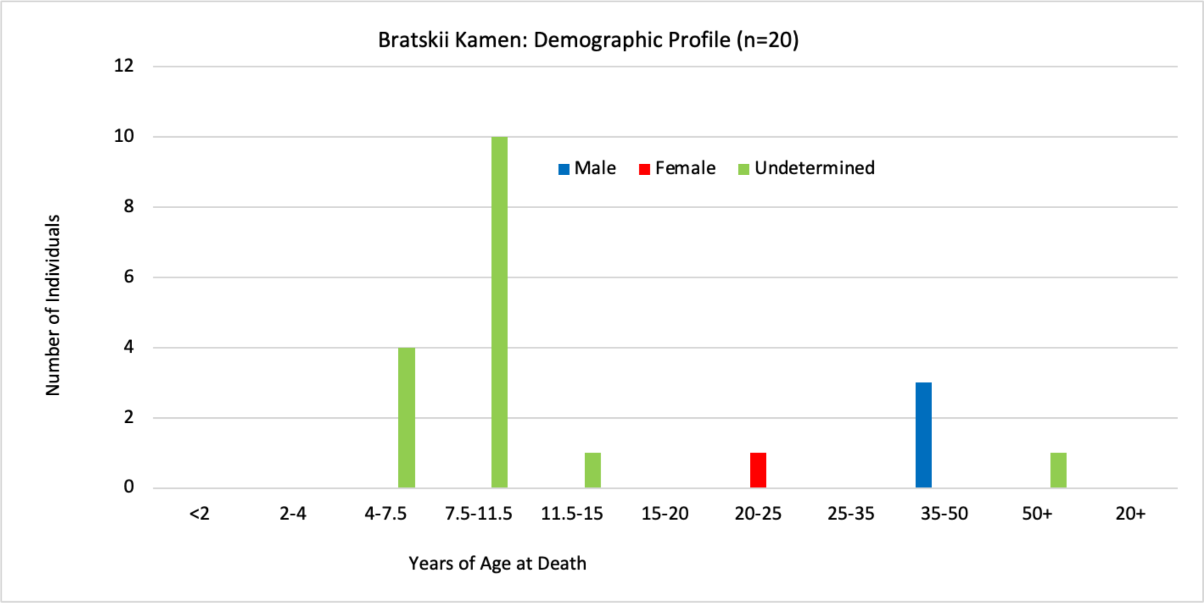


**Figure S4: Mortality profile inferred from age-at-death estimates for Late Neolithic individuals at Bratskii Kamen. Data for 12 of the 20 individuals provided by NN Mamonova.**

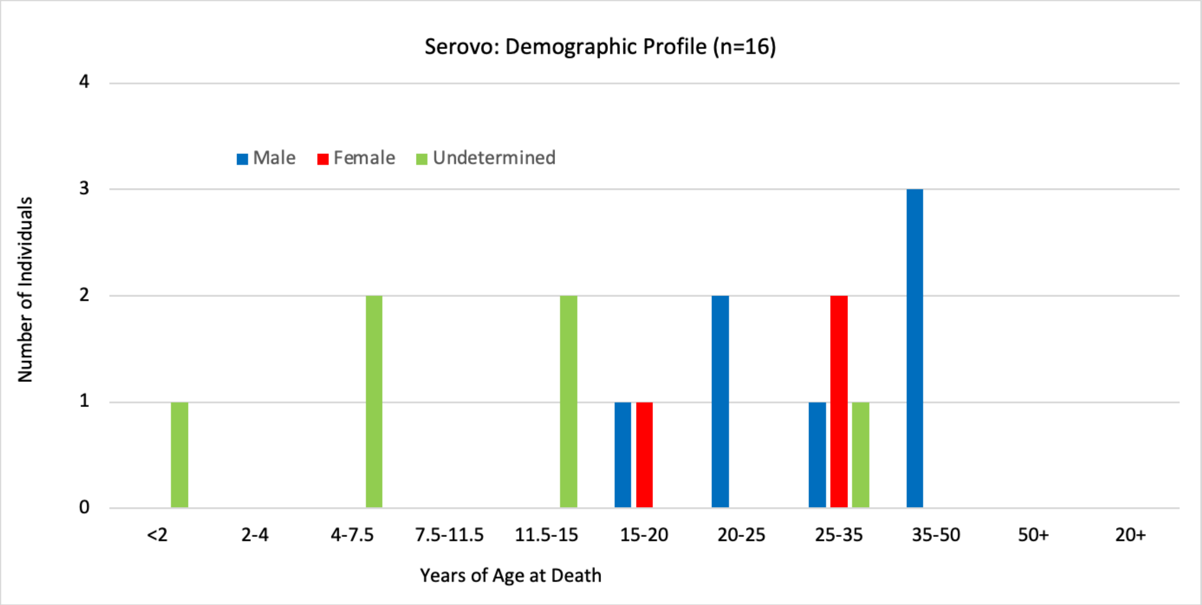


**Figure S5: Mortality profile inferred from age-at-death estimates for Late Neolithic individuals at Serovo. Data for 7 of the 16 individuals provided by NN Mamonova.**

### Supplementary Note 2: Ancient Human Genome Analysis

Uniparental Haplogroups

Uniparental haplogroups were assigned using mutserve and haplogrep for mitochondrial haplogroups as detailed above, and an in-house pipeline described in Seersholm et al.[20] for Y chromosome haplogroups. For the latter, individuals identified as XY genetic sex (*n*=22) with Y chromosome coverage >0.002 had Y chromosome-mapped reads SNP-called using bcftools mpileup and call, and the resulting genotypes used to assign haplogroups based on the corresponding derived alleles observed in ISOGG subgroups. Assigned haplogroups, quality scores and Y haplogroup paths are included in the Supplementary Data.

Y haplogroups were dominated by subgroups of Q1b~ (see Fig. S7), consistent with previous results showing the prevalence of this haplotype following the Middle Neolithic hiatus in the Cis-Baikal region[12] (Schulting et al. in prep.), Contrastingly, there is a much higher degree of mitochondrial haplotype diversity, even taking into account the higher sample size (*n*=46), an observation consistent with the scenario of male endogamy, and possible Y chromosomal bottleneck in the demographic history of this population.

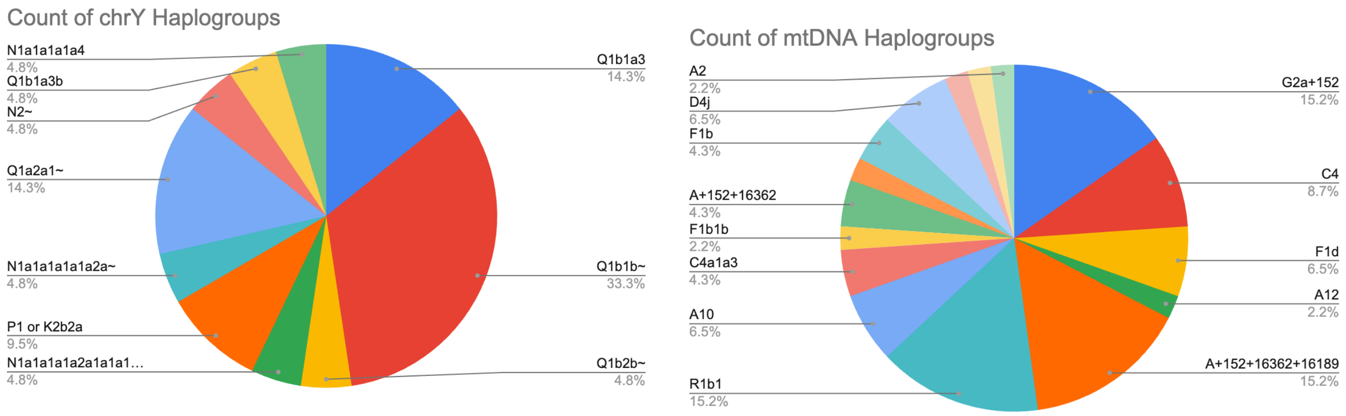


**Figure S7. Proportions of uniparental haplotypes observed in the 46 (24 XX, 22 XY) ancient individuals reported here.**

Inbreeding

Runs of Homozygosity (RoH) indicative of inbreeding were detected using two approaches: as Homozygous-by-Descent (HBD) segments reported by IBDseq, and with pseudohaploid genotypes from comparison with a large database of reference haplotypes using hapRoH. For hapRoH analysis, pseudohaploid genotypes were generated at the 1240K positions (autosomes only) using bcftools mpileup and call to create a VCF, and then vcf2eigenstrat from gdc (https://github.com/mathii/gdc). Samples with fewer than 400,000 SNPs represented were discarded from analysis. HapRoH primarily reports RoHs longer than 4cM, for which very few were detected across all the studied individuals (see Fig. S8). Only five individuals had ROH segments longer than 4cM: Ust'Ida #25.03 (CGG024154) with 9.6cM; Ust'Ida #22 (DA342) with 8.6cM; Ust'Ida #56.01 (DA345) with 4.5cM; and Ust'Ida #14 (DA355) with 4.1cM.  When the minimum RoH length threshold was reduced to 2cM, all individuals were found to have RoHs in their genome, supporting the interpretation that this result is indicative of an overall large effective population size. This was estimated using the maximum likelihood approach in hapROH and was found to be 18,219 individuals (95%CI: 9,445–42,062). HBD segments detected by IBDseq similarly showed a very low incidence of RoHs, with the large majority detected in the range of 0.5-1cM, and none individually longer than the 4-8cM range.

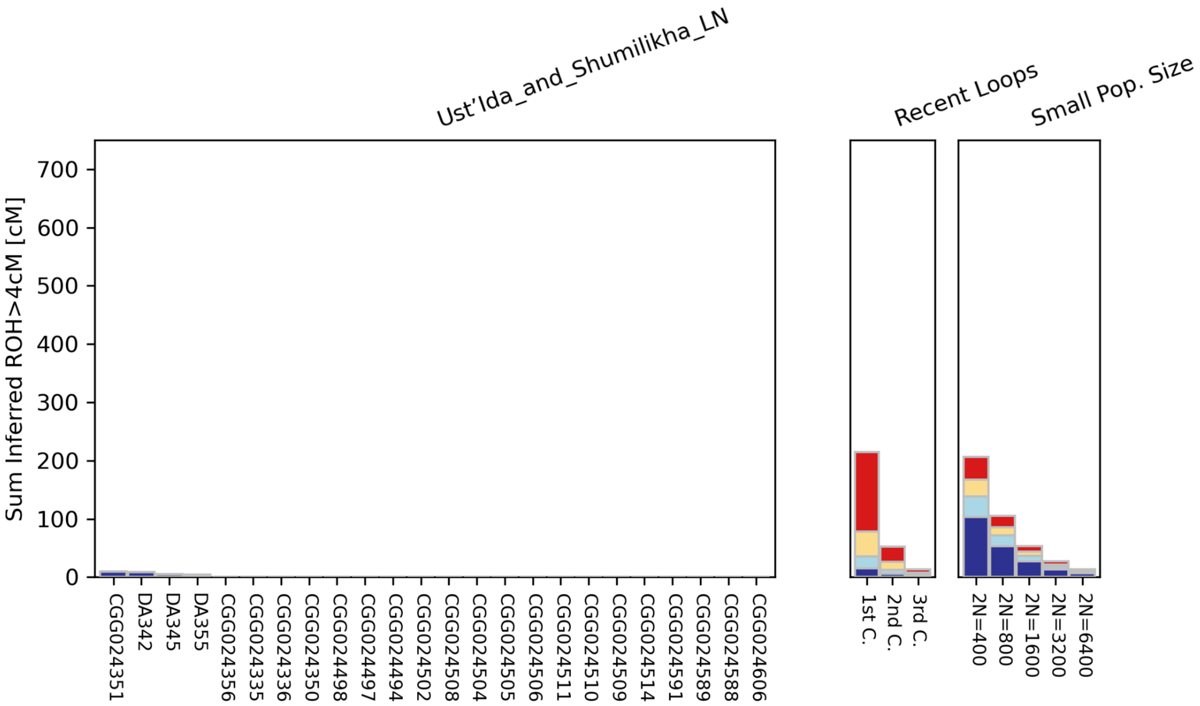


**Figure S8: Summed RoHs from Ust'-Ida and Shumilikha cemeteries, generated from hapRoH.** No RoHs >4cM were identified from individuals at Bratskii Kamen or Serovo.

<u>Kinship</u>

Pairwise biological kinship was inferred using KIN[21], which was found to have the highest reliability from a range of approaches previously tested in a Baikal hunter–gatherer population (Schulting et al. in prep.). Additionally, this approach is suitable for very low coverage data and was applied at the recommended minimum coverage of 0.05x. As such, the samples from Bratskii Kamen #17 (CGG024570) and Ust'-Ida I #26 (CGG024507) were omitted due to falling below this (0.006x and 0.01x respectively). This method applies a Hidden-Markov-Model approach to infer IBD fragments and estimate relatedness scenarios between pairs of individuals, returning log-likelihood estimates for the two most probable scenarios of relatedness. Pairwise kinship estimates and associated log-likelihoods are provided in Supplementary Data. Pedigrees were reconstructed on the following basis:

***Bratskii Kamen Pedigree (light blue)***: #19.01 (CGG024571) and #19.03 (CGG024573) are identified as third degree related; these share a grave and are inferred to have been buried contemporaneously at the same time (along with #19.02), and likely died contemporaneously; all are young girls. The ages of #19.01 and #19.03 are 8–9 years and 7–9 years respectively; these are therefore inferred as most likely cousins. #19.02 (CGG024572, 4–5 years) has autosomal coverage of 0.001, too low to infer kinship, but all three share the mitochondrial haplotype A+152+16362+16189, with three private mutations. As such, #19.01 and #19.03 are inferred to be maternal cousins, and #19.02 most likely related on their mothers' side.

***Shumilikha/Ust'-Ida I Pedigree (yellow)***: An avuncular relationship is identified between Shumilikha #2.01 (CGG024591) and Ust'-Ida #14 (DA355). This is inferred as paternal, as they have different mitochondrial haplogroups.

**Ust'-Ida Pedigrees:**
*(Blue)*: #8 (CGG024499), #26.01 (CGG024495) and #26.04 (CGG024510) are all identified as most likely full siblings of one another; all are of a similar age and share the mitochondrial haplogroup A10 (Y haplogroup could not be called for #8).

*(Green)*: #6 (CGG024350), 35–50 years, is identified as most likely the father of #33.01 (CGG024505), 12–25 years; they share Y haplotypes.

*(Red)*: #25.01 (CGG024514) and #25.02 (CGG024494) are full siblings (brother and sister) and share mitochondrial haplogroups; these are also 3rd/4th degree related to #6, and #25.01 shares Y haplotypes with him; as well as with #53.02 (CGG024501), 4–6 years, and has an avuncular relationship indicating #53.02 is a nephew on their brother's side.

*(Orange)*: #20.02 (CGG024335), 30–40 years, and #20.01 (CGG024336), 18–24 years; and #20.01 and #33.02 (CGG024502), 13–16 years, both have avuncular relationships, and ages and uniparental haplotypes that indicate they are the nephew and niece on a brother and sister's side respectively. #20.01 and #33.02 are also identified as most likely third degree related (cousins).

*(Brown)*: #18 (CGG0244498), 11–13 years, is the nephew of #56.01 (DA345), 35–50 years, on his father's side, on the basis of their avuncular relationship, haplogroups, and ages. #56.01 is also the grandfather of #44.01 (CGG024504), her father's father as they do not share mitochondrial haplogroups. In turn, #44.01 is the half-sibling of #44.02 (CGG024492) with a shared mother (having the same mitochondrial haplotypes), and she is also third degree related to DA345 (though not inbred).

| Sample ID 1 | Sample ID 2 | Relationship | Loglikelihood | 2nd Estimate | Loglikelihood 2nd Estimate |
|---|---|---|---|---|---|
| CGG024335 | CGG024336 | Avuncular | -1569.656107 | Half-siblings | -1569.817416 |
| CGG024335 | CGG024495 | 4th Degree | -1239.802955 | 5th Degree | -1240.613968 |
| CGG024335 | CGG024496 | 4th Degree | -1324.399894 | 5th Degree | -1324.533035 |
| CGG024335 | CGG024499 | 4th Degree | -1021.601223 | 3rd Degree | -1022.397917 |
| CGG024335 | CGG024502 | Avuncular | -1492.480266 | Half-siblings | -1493.397287 |
| CGG024335 | CGG024510 | 4th Degree | -1477.958189 | 3rd Degree | -1479.387116 |
| CGG024335 | CGG024606 | 4th Degree | -1408.265225 | 5th Degree | -1408.41702 |
| CGG024336 | CGG024502 | 3rd Degree | -1459.113319 | 4th Degree | -1460.496886 |
| CGG024350 | CGG024505 | Parent-Child | -1324.877862 | Grandparent-Grandchild | -1357.698779 |
| CGG024350 | CGG024514 | 4th Degree | -1435.728472 | 3rd Degree | -1436.575648 |
| CGG024351 | CGG024492 | 4th Degree | -1067.442709 | 5th Degree | -1067.820852 |
| CGG024351 | CGG024499 | 4th Degree | -1061.451395 | 5th Degree | -1061.571487 |

| | | | | | |
|---|---|---|---|---|---|
| CGG024351 | CGG024504 | 4th Degree | -1650.640775 | 3rd Degree | -1652.568132 |
| CGG024477 | CGG024577 | 4th Degree | -1517.45561 | 5th Degree | -1517.957534 |
| CGG024492 | CGG024504 | Half-siblings | -1032.903872 | Avuncular | -1033.562598 |
| CGG024492 | DA345 | 3rd Degree | -1047.420929 | 4th Degree | -1050.443868 |
| CGG024494 | CGG024501 | Avuncular | -1140.836206 | Half-siblings | -1141.09663 |
| CGG024494 | CGG024510 | 4th Degree | -1406.964352 | 5th Degree | -1407.40478 |
| CGG024494 | CGG024514 | Siblings | -1477.833312 | Grandparent-Grandchild | -1516.248223 |
| CGG024494 | DA355 | 4th Degree | -1358.339584 | 5th Degree | -1358.607971 |
| CGG024495 | CGG024496 | 4th Degree | -1040.908687 | 5th Degree | -1041.221649 |
| CGG024495 | CGG024499 | Siblings | -776.6951762 | Grandparent-Grandchild | -780.8381443 |
| CGG024495 | CGG024504 | 4th Degree | -1229.433845 | 5th Degree | -1230.499444 |
| CGG024495 | CGG024510 | Siblings | -1188.717673 | Grandparent-Grandchild | -1198.663965 |
| CGG024496 | CGG024499 | 4th Degree | -834.9894087 | 3rd Degree | -835.5851588 |
| CGG024496 | CGG024502 | 4th Degree | -1240.599549 | 5th Degree | -1241.177062 |
| CGG024496 | CGG024508 | 4th Degree | -1253.108589 | 5th Degree | -1253.710634 |
| CGG024496 | CGG024510 | 4th Degree | -1257.042477 | 5th Degree | -1258.562896 |
| CGG024496 | DA342 | 3rd Degree | -1442.464044 | 4th Degree | -1444.862556 |
| CGG024497 | CGG024498 | 4th Degree | -1352.245609 | 5th Degree | -1353.320999 |
| CGG024497 | CGG024589 | 4th Degree | -1199.801311 | 5th Degree | -1200.688314 |
| CGG024498 | CGG024504 | 4th Degree | -1548.445131 | 5th Degree | -1548.788624 |
| CGG024498 | CGG024508 | 4th Degree | -1461.810529 | 3rd Degree | -1461.964927 |
| CGG024498 | DA345 | Avuncular | -1583.41066 | Half-siblings | -1584.168584 |
| CGG024499 | CGG024501 | 4th Degree | -739.5163725 | 5th Degree | -740.4712195 |
| CGG024499 | CGG024502 | 4th Degree | -933.5957121 | 3rd Degree | -933.9783591 |
| CGG024499 | CGG024504 | 4th Degree | -1024.084358 | 5th Degree | -1024.126123 |
| CGG024499 | CGG024510 | Siblings | -957.1921583 | Grandparent-Grandchild | -966.029845 |
| CGG024500 | DA342 | 4th Degree | -1078.863973 | 5th Degree | -1079.215035 |
| CGG024501 | CGG024514 | 3rd Degree | -1112.972586 | 4th Degree | -1113.81473 |
| CGG024502 | CGG024508 | 4th Degree | -1365.11983 | 5th Degree | -1366.723952 |
| CGG024502 | CGG024510 | 3rd Degree | -1392.477855 | 4th Degree | -1394.166535 |
| CGG024502 | CGG024606 | 4th Degree | -1311.873669 | 3rd Degree | -1312.666053 |
| CGG024504 | CGG024510 | 4th Degree | -1457.756551 | 5th Degree | -1457.946806 |
| CGG024504 | DA345 | Grandparent-Grandchild | -1602.700396 | Half-siblings | -1603.179338 |
| CGG024504 | DA355 | 4th Degree | -1443.146164 | 3rd Degree | -1444.005127 |
| CGG024506 | DA344 | 3rd Degree | -1191.25556 | 4th Degree | -1194.068054 |

| | | | | | |
|---|---|---|---|---|---|
| CGG024508 | DA342 | 3rd Degree | -1599.341823 | 4th Degree | -1602.824445 |
| CGG024508 | DA344 | 4th Degree | -1209.271985 | 5th Degree | -1210.406751 |
| CGG024508 | DA345 | 4th Degree | -1492.292225 | 5th Degree | -1493.777996 |
| CGG024510 | DA342 | 4th Degree | -1565.04018 | 5th Degree | -1566.627835 |
| CGG024512 | DA344 | 4th Degree | -1167.71459 | 5th Degree | -1168.211886 |
| CGG024514 | DA345 | 4th Degree | -1480.631077 | 5th Degree | -1481.948266 |
| CGG024514 | DA355 | 4th Degree | -1324.419028 | 5th Degree | -1324.555557 |
| CGG024565 | DA355 | 4th Degree | -1184.714436 | 5th Degree | -1184.9387 |
| CGG024569 | CGG024574 | 4th Degree | -1244.008591 | 5th Degree | -1244.016551 |
| CGG024571 | CGG024573 | 3rd Degree | -950.5271721 | 4th Degree | -950.7023466 |
| CGG024571 | DA355 | 4th Degree | -1092.442289 | 5th Degree | -1092.799751 |
| CGG024576 | CGG024577 | 4th Degree | -1469.028639 | 5th Degree | -1470.64048 |
| CGG024589 | DA355 | 4th Degree | -1222.903369 | 5th Degree | -1223.973813 |
| CGG024591 | DA355 | Avuncular | -1293.3551 | 4th Degree | -1294.743272 |
| CGG024592 | CGG024598 | 3rd Degree | -800.6971728 | 4th Degree | -801.808945 |
| CGG024592 | DA355 | 4th Degree | -1055.153991 | 5th Degree | -1055.958448 |
| DA342 | DA344 | 4th Degree | -1327.884711 | 5th Degree | -1327.893621 |

**Table S1: Pairwise kinship estimates from KIN, selecting only those at least 4th degree in the most likely resulting relationship scenario.**

Principal components Analysis

Ancestry of ancient individuals was explored by Principal Components Analysis (PCA) as detailed in the Methods. This showed that all individuals clustered as expected with previously published genomes from Late Neolithic and Early Bronze Age Baikal hunter–gatherers, at the 'East' end of the Eastern/Western Hunter–Gatherer ancestry cline (Fig. S9.)

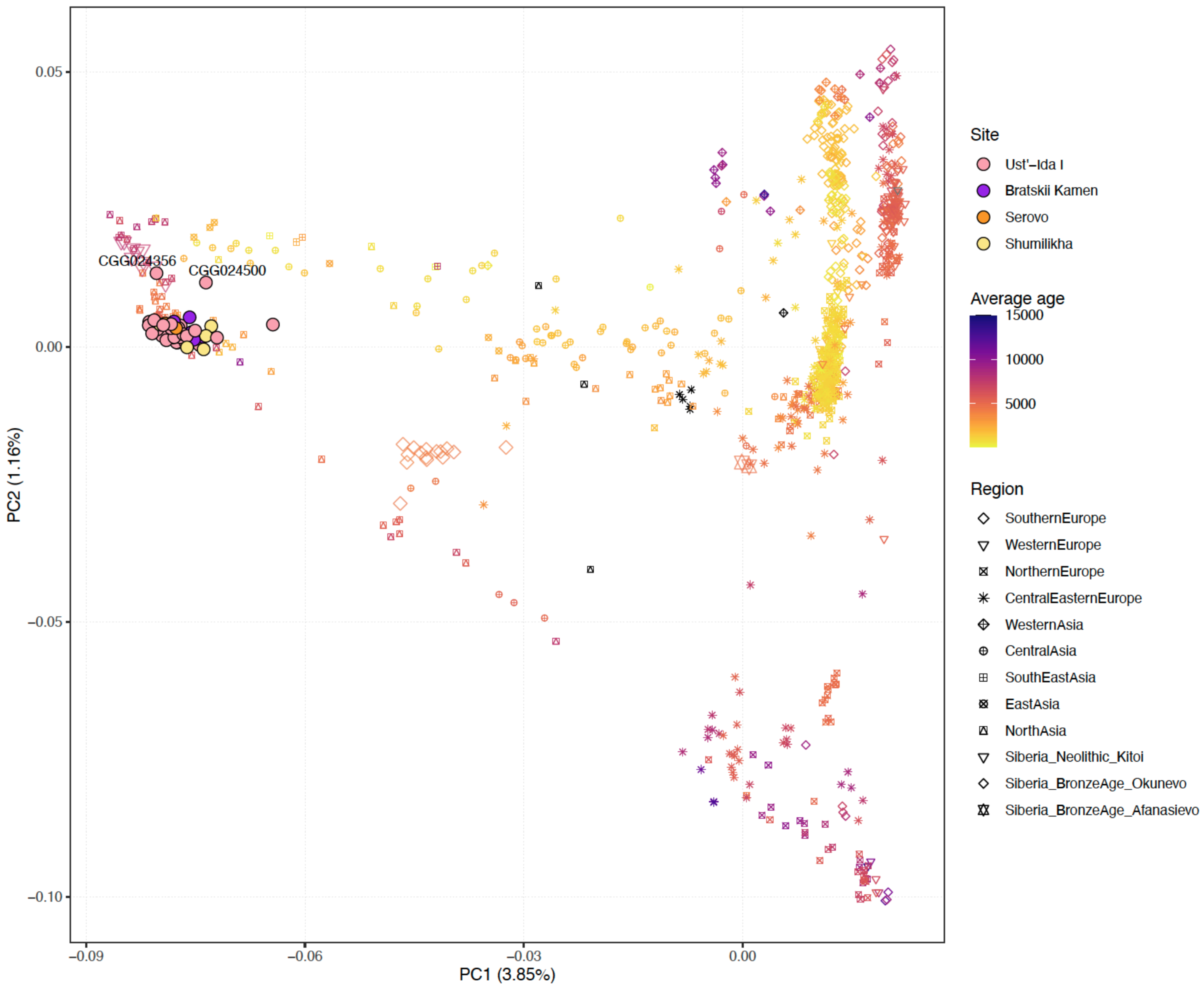


**Figure S9.** Principal Components Analysis of genomes for 46 ancient individuals studied here (31 Ust'-Ida I; 8 Bratskii Kamen; 2 Serovo; and 5 Shumilkha).

#### Supplementary Note 3: Ancient Pathogen DNA Analysis

Metagenomic detection of Pathogens

Following filtering of pairwise alignment results with the thresholds described in Methods, the most predominant microbial pathogen identified among shotgun-sequenced ancient DNA data was *Yersinia pestis*, with 525,355 uniquely aligned reads across screening data from all samples (see Supplementary Data). Higher coverage plague genomes were utilised for downstream analysis, following merging with genome capture data where this was performed (Table S2). Aside from *Y. pestis* taxonomic identifications, the most frequently identified bacteria are of *Pseudomonas* spp. (from 22 individuals), followed by *Streptococcus spp.* and *Acinetobacter spp.* (from 15 individuals each). In most of these cases, it appears that these can be attributed to environmental strains of these taxa, for example Acinetobacter sp. DUT-2, where reference data were sequenced from marine sediments (https://www.ncbi.nlm.nih.gov/biosample/SAMN04525297). Nonetheless, it is potentially significant that overall, 74% of the identified microbes within these ancient samples are gram-negative. Though many of the non-environmental gram negative bacteria are harmless to humans (e.g. *Wolbachia sp.* and *Brevibacterium flavum*) or are oral symbionts (e.g. *Streptococcus vestibularis*), a number of these are of note as reported human pathogens.

The significance of the identification of *Streptococcus pseudopneumoniae* (identified in Bratskii Kamen #19.01, CGG024571) is also unclear; while it has been suggested this may have a clinical significance in pneumonia infection, this remains under debate[22,23]. It is important to note that pathogenesis is not only a microbial trait but an outcome of the interplay between a host and a microbe. While it is likely that *Y. pestis* caused severe disease in all those infected, the consequences of the presence of many other microbes, especially those sometimes termed opportunistic pathogens, are more challenging to appreciate. Today, we find that while some microbes, e.g. *Streptococcus pseudopneumoniae,* preferentially cause disease in those with weakened immune systems, they can also sometimes cause disease in those with healthy immune systems. Likewise, the immune system's efficiency can be affected by starvation or malnutrition in those not otherwise immunocompromised; this might facilitate serious infections by microbes that otherwise would not have been a likely cause of disease. In some cases, the host's immune response to infection causes the most severe symptoms, e.g., in infectious mononucleosis (primarily caused by Epstein-Barr virus (EBV) infection in teenagers), sepsis, or toxic shock syndrome, typically caused by some gram-positive bacteria or *Y. pseudotuberculosis*). Bacterial superinfections following viral infections or bacterial co-infections further complicate host susceptibility to a severe outcome. Moreover, the pathogenic potential of specific microbes in the past might not fully be reflected in the most similar variants today. Consequently, the consequences of many of the microbes found in aDNA samples are challenging to appreciate fully.

| **sampleId** | **Site (burial number)** | **Shotgun coverage** | **merged capture and shotgun coverage** |
|---|---|---|---|
| CGG024508 | Ust'-Ida I (#31) | 0.174 | 0.659 |
| CGG024510 | Ust'-Ida I (#26.0) | 0.073 | - |
| CGG024511 | Ust'-Ida I (#32) | 0.095 | - |
| CGG024571 | Bratskii Kamen (#19.01) | 0.06 | - |
| CGG024572 | Bratskii Kamen (#19.02) | 0.03 | - |
| CGG024573 | Bratskii Kamen (#19.03) | 0.116 | - |
| CGG024574 | Bratskii Kamen (#22) | 0.742 | 1.577 |
| CGG024576 | Serovo (#10) | 0.935 | 0.976 |
| CGG024588 | Ust'-Ida I (#16.01) | 0.304 | 0.733 |
| CGG024606 | Shumilikha (#34) | 2.619 | 6.358 |

**Table S2. Coverages for plague genomes with sufficient data to be included in the phylogeny (Fig. 2).**

Brucellosis in a Prehistoric Hunter–gatherer

The identification of *Brucella sp.* in Ust'Ida I #26.04 (CGG024495) is the first detection of this pathogen to our knowledge in a prehistoric human. From the *pathopipe* pipeline[24], this was identified as *B. canis*, the causative agent of canine brucellosis, a highly contagious infection in dogs and other canids transmitted through sexual contact or bodily fluids[25]. This identification is based on 96 uniquely aligned reads (covering 3009 bps in total) for this bacterium, passing all the additional filtering thresholds described above. While there are relatively few reads from which to estimate DNA damage patterns, we observe that the damage associated misincorporation rate at the 3' end of strands is 5.26%. Given the very small amount of genome coverage available, and high sequence similarity between *B. canis* and other *Brucella* species[26], we are unable to report with certainty that it is this specific species.

Nonetheless, of the principle species of animal hosts known to transmit the disease to humans today (cattle, buffalo, goats, sheep, camel, pigs and dogs[26]), only dogs (and other canids) are known from archaeological assemblages at Lake Baikal at this point in time. Infection can lead to infertility in both sexes, as well as, e.g., anorexia, pain, and lameness. Dog-to-human transmission is relatively rare and today usually contingent on a broken skin barrier and contact with gentital fluids from a birthing dog or a dog who has had a miscarriage. Human infection results in a febrile syndrome characterised by general symptoms such as splenomegaly, fatigue, and weakness[27]. The appearance of symptoms from infection can be delayed from weeks to years, and transmission of infection between humans is extremely rare. As such, infection in this individual would almost certainly have been the result of direct zoonotic transfer from an infected canid, and would have been very unlikely to have been the cause of death (at least in itself). The affected individual is a 10–12 years old male, directly dated to 5585–5065 cal. BP, and who is buried alongside his elder sister (positive for plague) in the shared grave 26, as well as having a brother located in the nearby grave number 8. This individual was additionally noted as slightly below the threshold number of reads for confident identification of *Y. pestis*.

Dogs are well-attested archaeologically in association with mid-Holocene hunter–gatherers at Lake Baikal, and dog burials in human-like graves occurred in Early Neolithic *Kitoi* cemetery sites, though no canid remains have been identified as dating to the Late Neolithic period at Baikal so far[28]. The mortuary treatment of dogs during this preceding period nonetheless implies the attachment of significant cultural importance to human relationships with these animals, whether symbolic or practical[29]. Additionally, wolves (*Canis lupus*) are known from burial sites in the Early Neolithic (and even a bear burial at Shamanka II), and although stable isotope analysis has suggested that dogs were eating a similar diet to humans[28], it is unclear as to the extent dogs filled a similarly undomesticated niche to wolves at this point in time or not. Infection could therefore have come equally from either a dog within a commensal or domesticate context of association with humans, or a wild dog or wolf. Consumption of canid remains, or the ritual exposure of bodily fluids (particularly in relation to the sexual organs) might explain this finding, especially in the context of earlier archaeological evidence for ritualised treatment of canid remains. Overall, this result further emphasises the significant role which zoonotic pathogens likely played in the lives of prehistoric hunter–gatherers.

#### Phylogenetic Modelling and Bayesian Estimation of Node Dates

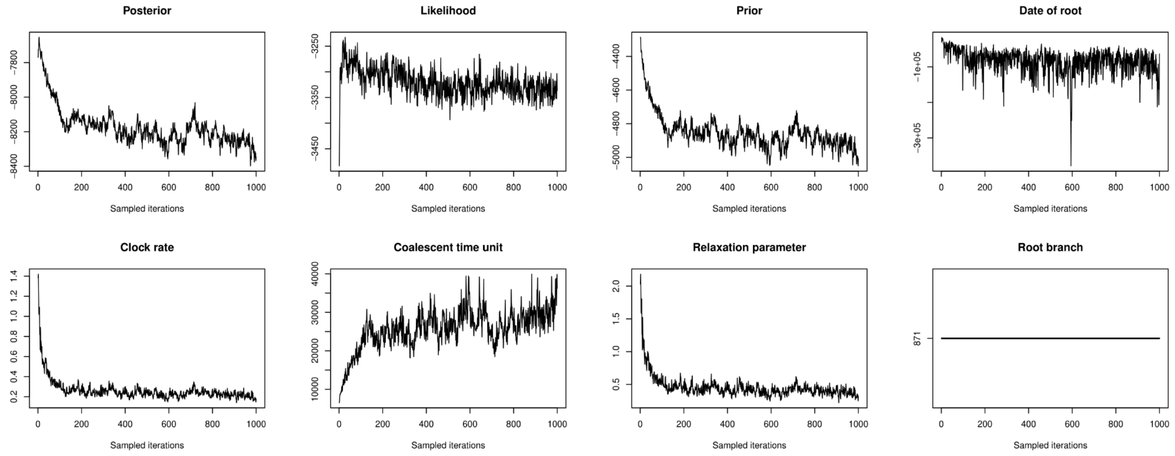


**Figure S10: The trace from BactDating after 1 million iterations.** This shows convergence for the estimate of molecular divergence dates. Iterations were sampled every 1000 iterations.

The newick file for the full phylogenetic tree created with RAxML is available on the Github, as is the json file for the USHER tree annotated with substitution information which can be uploaded to auspice.us to obtain an interactive phylogeny.

#### Classic *Y. pestis* Virulence Genes

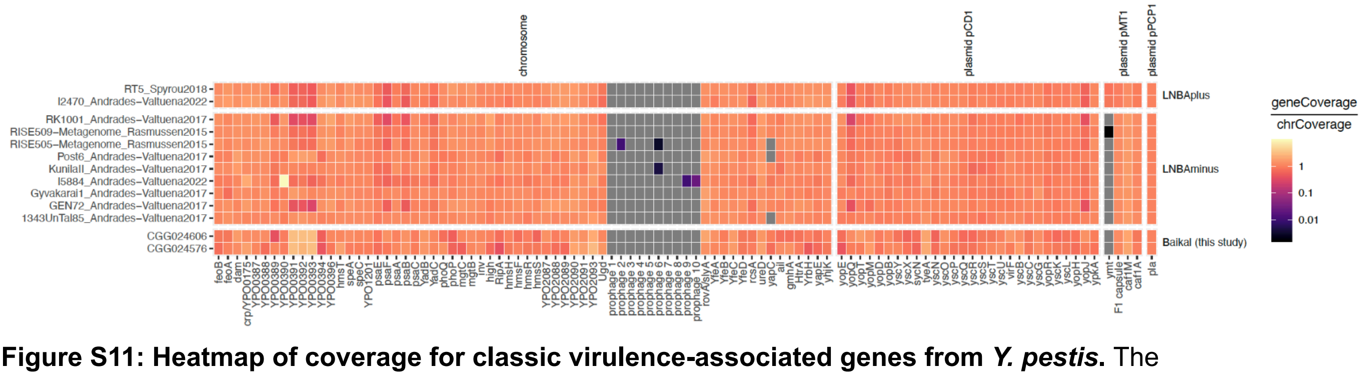


**Figure S11: Heatmap of coverage for classic virulence-associated genes from *Y. pestis*.** The two highest coverage Baikal samples, Shumilikha #34 (CGG024606) and Serovo #10 (CGG024576) are shown at the bottom alongside previously LNBA- and LNBA+ strains.

### Supplementary Note 4: Radiocarbon Chronology

Radiocarbon dating was undertaken through direct sampling of skeletal elements (principally either cranial, femoral or costal bone, see Supplementary Data). All subsampled material was processed at the Oxford Radiocarbon Accelerator Unit (ORAU) at the University of Oxford, UK. Carbon isotope measurements were made on ultra-filtered collagen following Brock et al. [30]. Additionally, collagen quality was assessed following the criteria of DeNiro [31] and van Klinken [32], with none of the reported samples rejected on the basis of these.

A total of 58 radiocarbon dates were used in modelled date ranges related to the outbreaks of plague described in this study, shown in Fig 1b. Of these, 22 radiocarbon dates are reported here for the first time; these comprise 10 from Late Neolithic individuals from Bratskii Kamen, 6 from Serovo, and 6 from Shumilikha. Raw dates, calibrated date ranges and metadata are provided for these samples, alongside the same for the 36 Late Neolithic individuals from Ust-'Ida I reported by Weber et al.[31], Weber et al.[33], Bronk Ramsey et al.[8]. To mitigate the Freshwater Reservoir Effect (FRE) resulting from human consumption of aquatic resources and metabolisation of older carbon, a correction was applied using the equation below, after Weber et al.[33], based on the regression model developed by Schulting et al.[34].

$$Y = -1388.8522 + 125.4503 \times \delta^{15}\mathrm{N}$$

**Equation 1.** Regression equation for the correction of radiocarbon dates from human skeletal tissue in mitigation of the Freshwater Reservoir Effect in the Angara Valley region of Cis-Baikal.

Radiocarbon dates were calibrated in OxCal v4.4[35] using atmospheric data from the IntCal20 calibration curve[36]. A Kernel Density Estimate function ("KDE_Model") [37] was used to model radiocarbon date ranges in OxCal. The resulting modelled dates are provided in Fig. S10. Additionally, OxCal code is provided below for this.

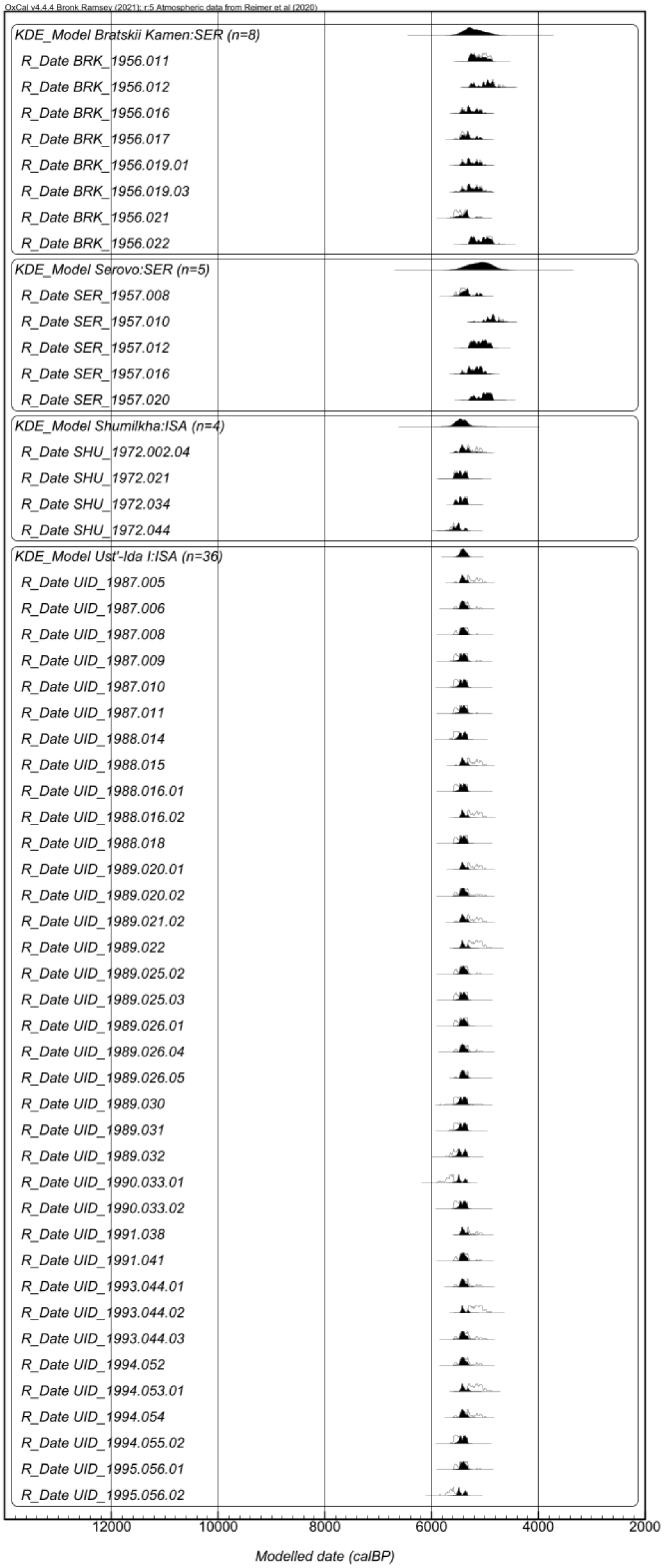

OxCal v4.4.4 Bronk Ramsey (2021); r:5 Atmospheric data from Reimer et al (2020)
KDE_Model Bratskii Kamen:SER (n=8)
R_Date BRK_1956.011
R_Date BRK_1956.012
R_Date BRK_1956.016
R_Date BRK_1956.017
R_Date BRK_1956.019.01
R_Date BRK_1956.019.03
R_Date BRK_1956.021
R_Date BRK_1956.022
KDE_Model Serovo:SER (n=5)
R_Date SER_1957.008
R_Date SER_1957.010
R_Date SER_1957.012
R_Date SER_1957.016
R_Date SER_1957.020
KDE_Model Shumilkha:ISA (n=4)
R_Date SHU_1972.002.04
R_Date SHU_1972.021
R_Date SHU_1972.034
R_Date SHU_1972.044
KDE_Model Ust'-Ida I:ISA (n=36)
R_Date UID_1987.005
R_Date UID_1987.006
R_Date UID_1987.008
R_Date UID_1987.009
R_Date UID_1987.010
R_Date UID_1987.011
R_Date UID_1988.014
R_Date UID_1988.015
R_Date UID_1988.016.01
R_Date UID_1988.016.02
R_Date UID_1988.018
R_Date UID_1989.020.01
R_Date UID_1989.020.02
R_Date UID_1989.021.02
R_Date UID_1989.022
R_Date UID_1989.025.02
R_Date UID_1989.025.03
R_Date UID_1989.026.01
R_Date UID_1989.026.04
R_Date UID_1989.026.05
R_Date UID_1989.030
R_Date UID_1989.031
R_Date UID_1989.032
R_Date UID_1990.033.01
R_Date UID_1990.033.02
R_Date UID_1991.038
R_Date UID_1991.041
R_Date UID_1993.044.01
R_Date UID_1993.044.02
R_Date UID_1993.044.03
R_Date UID_1994.052
R_Date UID_1994.053.01
R_Date UID_1994.054
R_Date UID_1994.055.02
R_Date UID_1995.056.01
R_Date UID_1995.056.02
12000
10000
8000
6000
4000
2000
Modelled date (calBP)

**Figure S12. Radiocarbon Date Probability Density Functions generated from the KDE_Model function in OxCal for Late Neolithic individuals at Bratskii Kamen, Serovo, Shumilikha and Serovo.**

#### OxCal Code

```
Plot()
 {
  KDE_Model("Bratskii Kamen:SER (n=8)")
  {
   MortTrad="Serovo";
   longitude=101.675;
   latitude=56.122222;
   R_DATE("BRK_1956.011",4443,68);
   R_DATE("BRK_1956.012",4287,68);
   R_DATE("BRK_1956.016",4586,68);
   R_DATE("BRK_1956.017",4623,68);
   R_DATE("BRK_1956.019.01",4580,68);
   R_DATE("BRK_1956.019.03",4565,68);
   R_DATE("BRK_1956.021",4734,68);
   R_DATE("BRK_1956.022",4372,70);
  };
  KDE_Model("Serovo:SER (n=5)")
  {
   MortTrad="SEROVO";
   longitude=103.222222;
   latitude=54.509722;
   R_DATE("SER_1957.008",4662,68);
   R_DATE("SER_1957.010",4263,66);
   R_DATE("SER_1957.012",4446,68);
   R_DATE("SER_1957.016",4550,69);
   R_DATE("SER_1957.020",4386,67);
  };
  KDE_Model("Shumilkha:ISA (n=4)")
  {
   MortTrad="Isakovo";
   longitude=101.675;
   latitude=52.906719;
   R_DATE("SHU_1972.002.04",4591,67);
   R_DATE("SHU_1972.021",4733,67);
   R_DATE("SHU_1972.034",4705,48);
   R_DATE("SHU_1972.044",4838,68);
  };
  KDE_Model("Ust'-Ida I:ISA (n=36)")
  {
   MortTrad="Isakovo";
   longitude=103.377078;
   latitude=53.182078;
```

```
    R_DATE("UID_1987.005",4596,73);
    R_DATE("UID_1987.006",4641,71);
    R_DATE("UID_1987.008",4666,72);
    R_DATE("UID_1987.009",4719,71);
    R_DATE("UID_1987.010",4743,71);
    R_DATE("UID_1987.011",4701,70);
    R_DATE("UID_1988.014",4774,71);
    R_DATE("UID_1988.015",4585,71);
    R_DATE("UID_1988.016.01",4730,70);
    R_DATE("UID_1988.016.02",4572,71);
    R_DATE("UID_1988.018",4710,71);
    R_DATE("UID_1989.020.01",4584,71);
    R_DATE("UID_1989.020.02",4655,75);
    R_DATE("UID_1989.021.02",4597,70);
    R_DATE("UID_1989.022",4543,71);
    R_DATE("UID_1989.025.02",4676,71);
    R_DATE("UID_1989.025.03",4710,71);
    R_DATE("UID_1989.026.01",4696,71);
    R_DATE("UID_1989.026.04",4646,72);
    R_DATE("UID_1989.026.05",4646,52);
    R_DATE("UID_1989.030",4750,72);
    R_DATE("UID_1989.031",4753,68);
    R_DATE("UID_1989.032",4826,71);
    R_DATE("UID_1990.033.01",4901,71);
    R_DATE("UID_1990.033.02",4741,72);
    R_DATE("UID_1991.038",4602,56);
    R_DATE("UID_1991.041",4663,72);
    R_DATE("UID_1993.044.01",4633,71);
    R_DATE("UID_1993.044.02",4530,71);
    R_DATE("UID_1993.044.03",4638,72);
    R_DATE("UID_1994.052",4644,71);
    R_DATE("UID_1994.053.01",4548,71);
    R_DATE("UID_1994.054",4617,74);
    R_DATE("UID_1994.055.02",4754,72);
    R_DATE("UID_1995.056.01",4680,73);
    R_DATE("UID_1995.056.02",4876,73);
  };
 };
```

#### Supplementary Note 5: Mortality Profile Modelling

<u>Analysis of Age-at-death Profiles</u>

We tested two hypotheses. Firstly that plague phases had a lower than expected average age-at-death. Secondly, that plague phases had an unusually high proportion of juvenile deaths (8-12 yrs old), given the assumption that this specific age range was more susceptible.

Stage 1 requires building a null model for the continuous probability of death at any age. We excluded four individuals who died unborn in-utero. Four further individuals were reported with the false precision of a point age (p) rather than a range. These were given a range of 0.8p to 1.2(1/6 +p) where p was measured in years. This resulted in a dataset of 625 individuals with osteometric age ranges.

Preliminarily, several continuous parametric age-at-death models were considered. The best fitting was a two-mixture model of a half-gaussian (encompassing the high proportion of deaths expected soon after birth) and a truncated gaussian (encompassing deaths into normal adulthood). This model required four parameters: the SD of the half-gaussian, the mean and SD of the truncated gaussian, and the mixture proportion of the half-gaussian. Maximum Likelihood parameters were found using the JDEoptim function from the R package DEoptimR[38]. This was achieved by calculating the model PDF across the osteometric age range of each individual.

Phases were selected with n = 20 or more individuals, providing 9 phases to test. This was achieved by comparing the two summary statistics of interest (proportion of juvenile deaths in each phase, and mean age-at-death), with the same summary statistics when sampling n random individuals from the null distribution, where n is the total number of deaths in the test phase. This is conservative test, since the deaths in the test phase always contribute to the null distribution. Significance was calculated as follows.

To calculate summary statistics from the phase of interest:

1. Each individual was assigned a point age, by randomly sampling from the null across their osteometric age ranges.
2. Summary statistics were calculated.
3. Steps 1 and 2 were repeated 5000 times to give a distribution of each summary statistic.

To calculate summary statistics from the null distribution:

1. We sampled n individuals from the full dataset, where n is the number of individuals in the test phase.
2. Each individual was assigned a point age, by randomly sampling from the null across their osteometric age range.
3. Summary statistics were calculated.
4. Steps 1 to 3 were repeated 5000 times.

A one-tailed p-value for the mean age (lower than expected) was calculated by fitting a gaussian to the null distribution of mean ages, and calculating the proportion of the PDF below the observed mean age. This was averaged across all 5000 resamplings.

A one-tailed p-value for the proportion of juveniles (higher than expected) was calculated by fitting a binomial to the mean proportion of null juveniles, then calculating the proportion of the PDF above the observed proportion of juveniles. This was averaged across all 5000 resamplings.

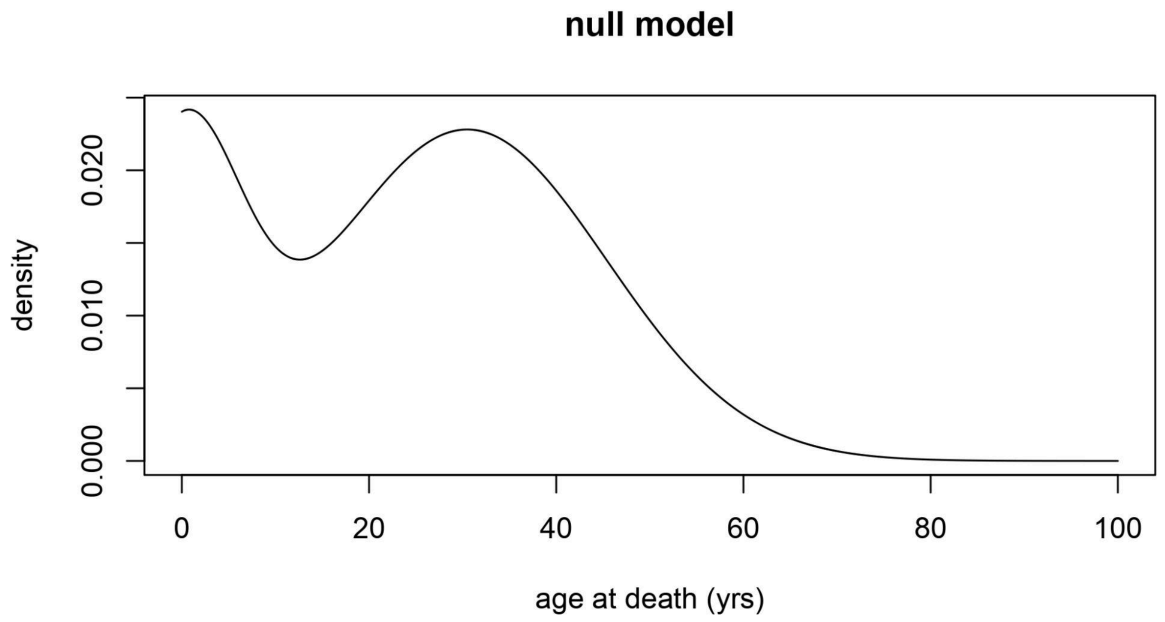


**Figure S13. Continuous null model of age-at-death.** Fitted to n=625 individuals with osteometric age ranges. Best fitting model was a two-mixture model of a half-gaussian (SD = 0.201) and a truncated Gaussian (mean = 30.492, SD =14.883, zero probability for age<=0), with the half-gaussian comprising 16.63% of the mixture.

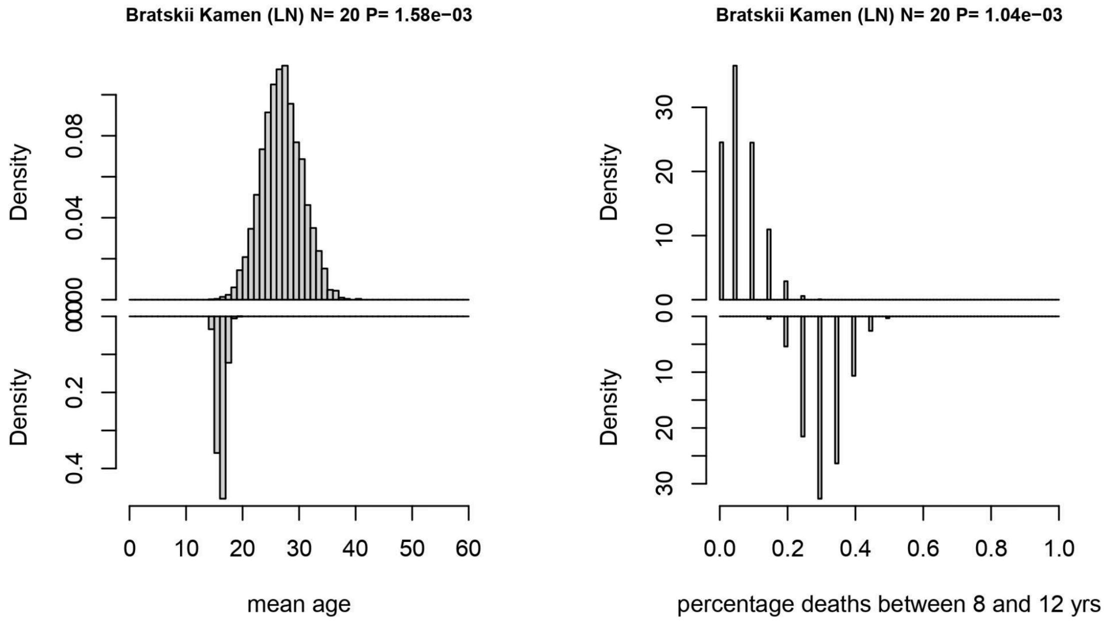


**Figure S14. Age-at-death modelling for Bratskii Kamen (LN).** Left: null distribution of mean age (top), test phase distribution of mean age (bottom). Right: null distribution of percentage of juvenile percentage (top), test phase distribution of juvenile percentage (bottom).

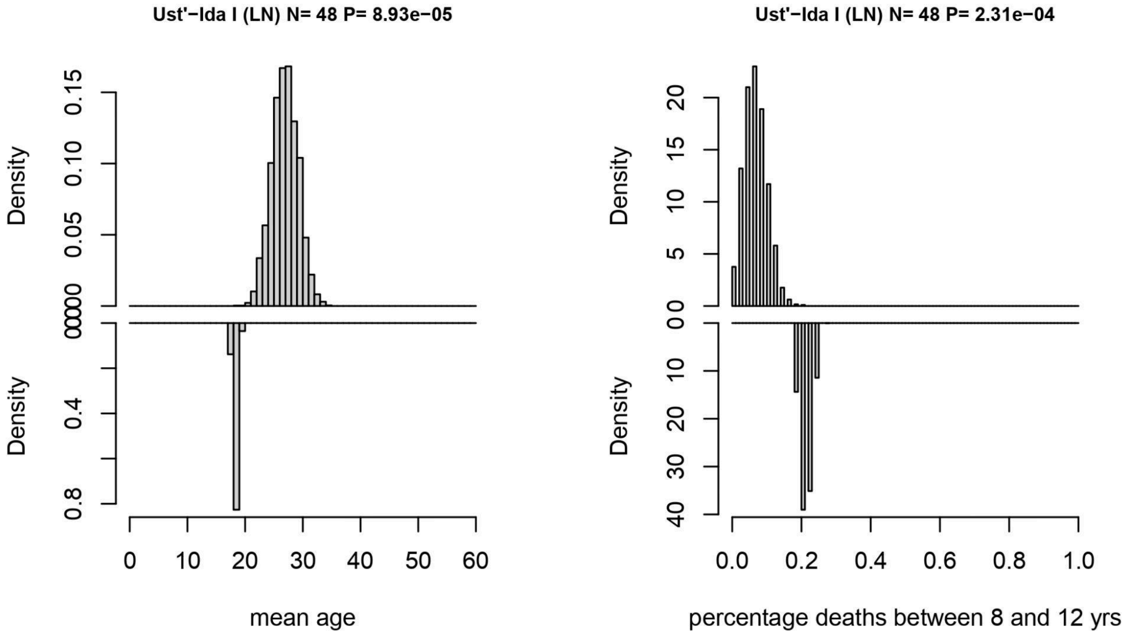


**Figure S15. Age-at-death modelling for Ust-Ida I (LN).** Left: null distribution of mean age (top), test phase distribution of mean age (bottom). Right: null distribution of percentage of juvenile percentage (top), test phase distribution of juvenile percentage (bottom).

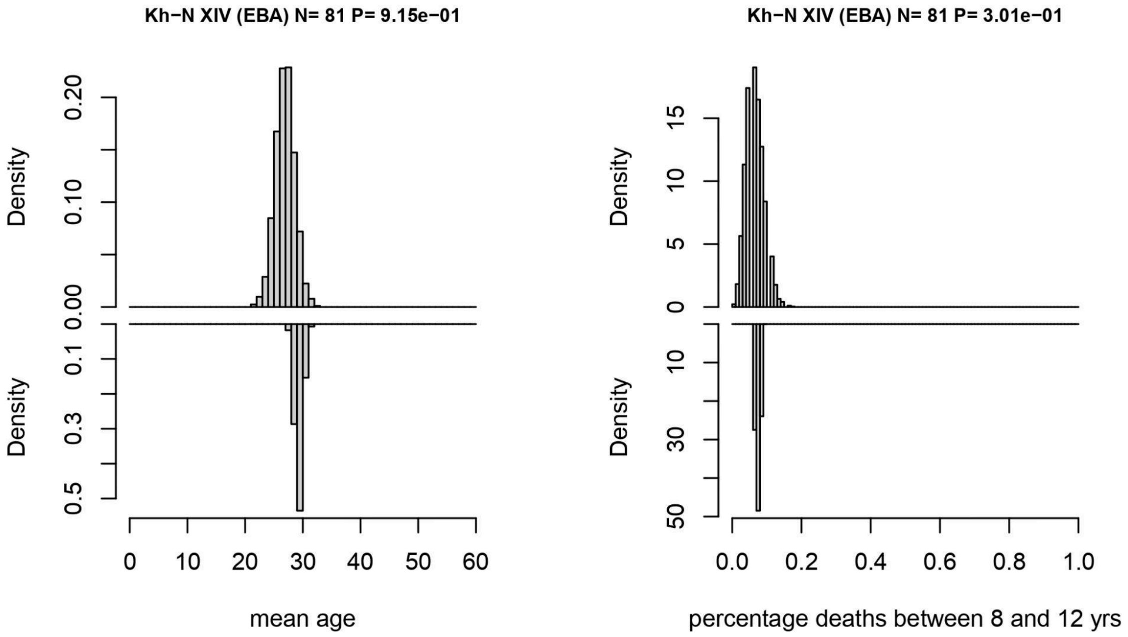


**Figure S16. Age-at-death modelling for Kuzhir-Nuge XIV.** Left: null distribution of mean age (top), test phase distribution of mean age (bottom). Right: null distribution of percentage of juvenile percentage (top), test phase distribution of juvenile percentage (bottom).

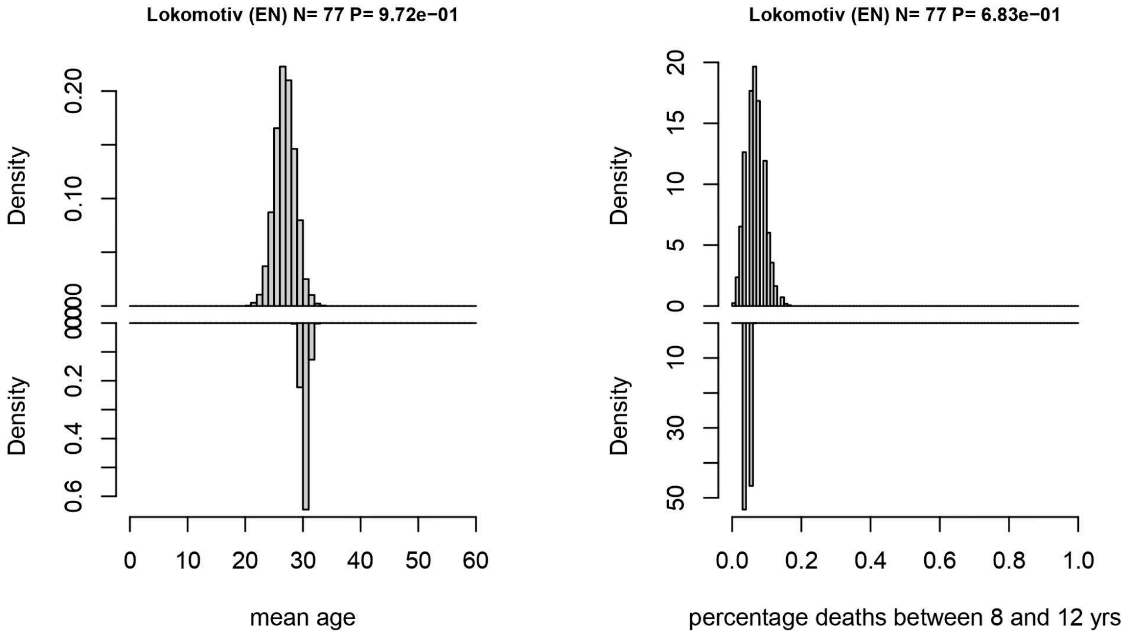


**Figure S17. Age-at-death modelling for Lokomotiv.** Left: null distribution of mean age (top), test phase distribution of mean age (bottom). Right: null distribution of percentage of juvenile percentage (top), test phase distribution of juvenile percentage (bottom).

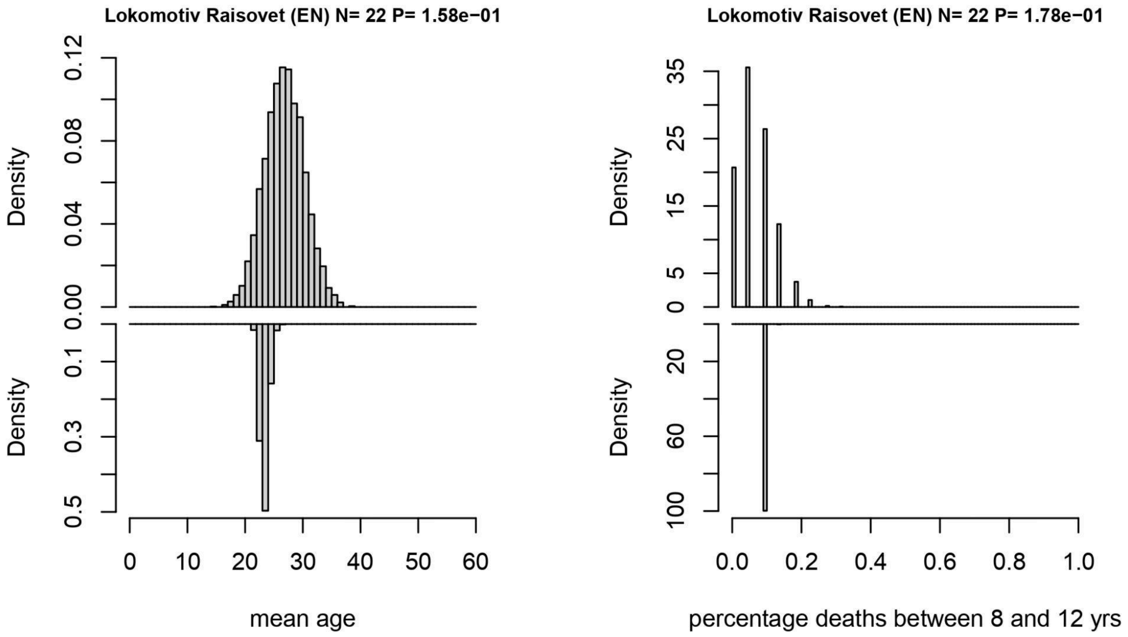


**Figure S18. Age-at-death modelling for Lokomotiv Raisovet.** Left: null distribution of mean age (top), test phase distribution of mean age (bottom). Right: null distribution of percentage of juvenile percentage (top), test phase distribution of juvenile percentage (bottom).

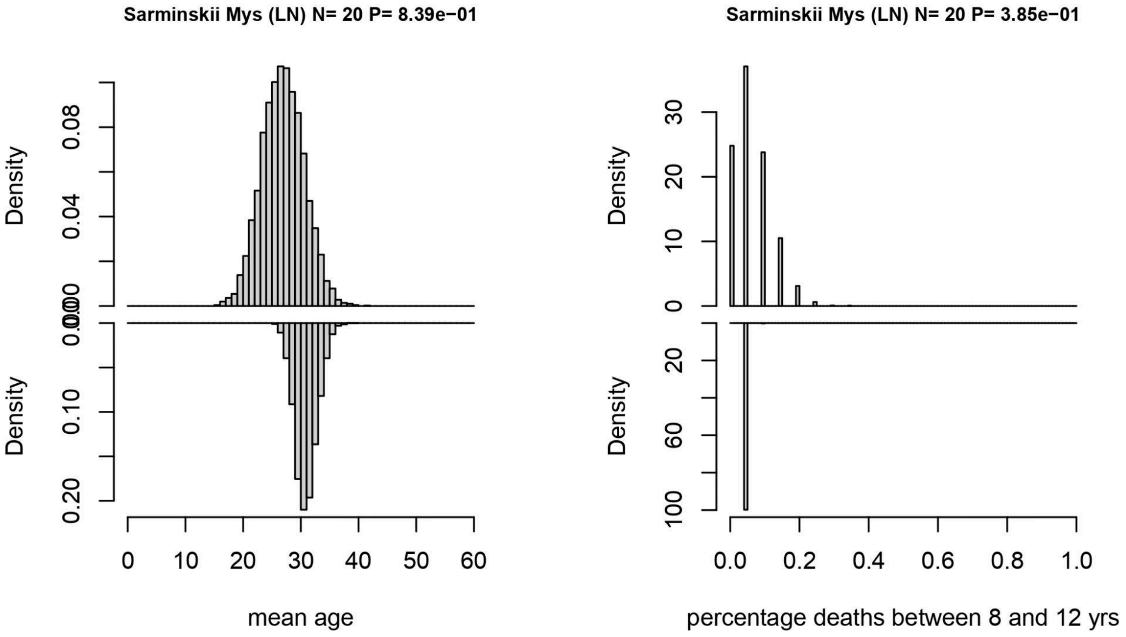


**Figure S19. Age-at-death modelling for Sarminskii Mys.** Left: null distribution of mean age (top), test phase distribution of mean age (bottom). Right: null distribution of percentage of juvenile percentage (top), test phase distribution of juvenile percentage (bottom).

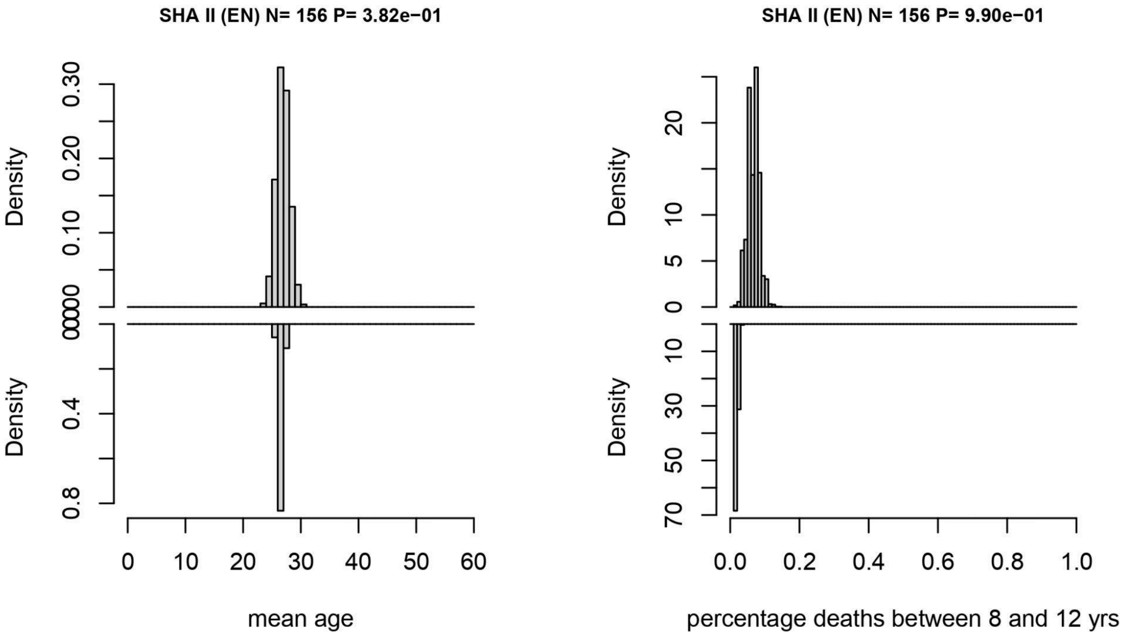


**Figure S20. Age-at-death modelling for Shamanka II.** Left: null distribution of mean age (top), test phase distribution of mean age (bottom). Right: null distribution of percentage of juvenile percentage (top), test phase distribution of juvenile percentage (bottom).

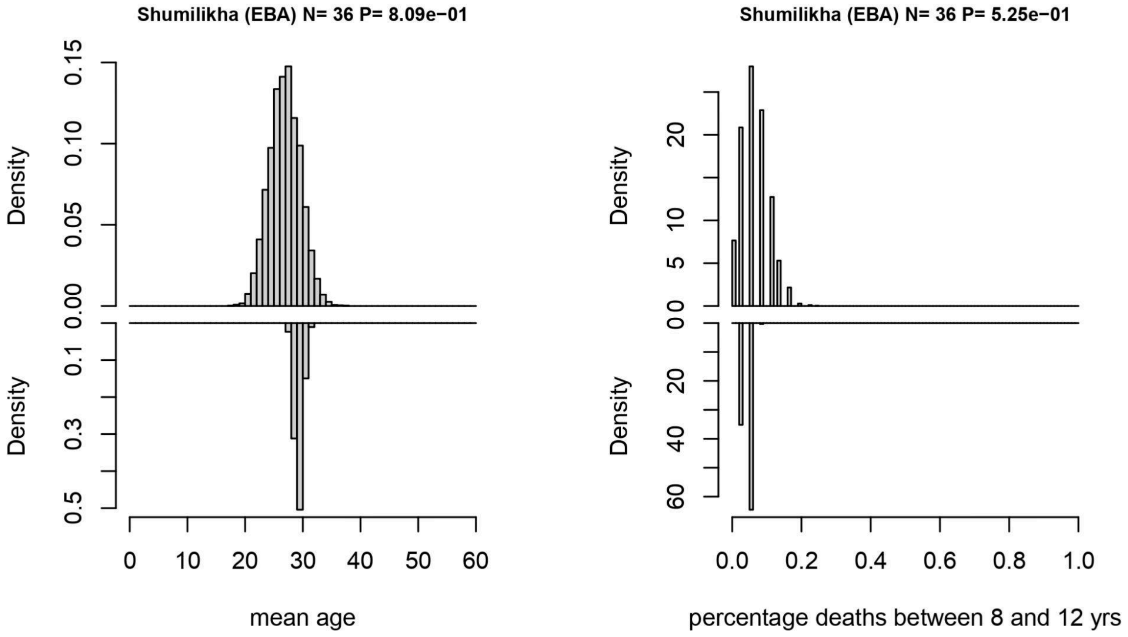


**Figure S21. Age-at-death modelling for Shamanka II.** Left: null distribution of mean age (top), test phase distribution of mean age (bottom). Right: null distribution of percentage of juvenile percentage (top), test phase distribution of juvenile percentage (bottom).

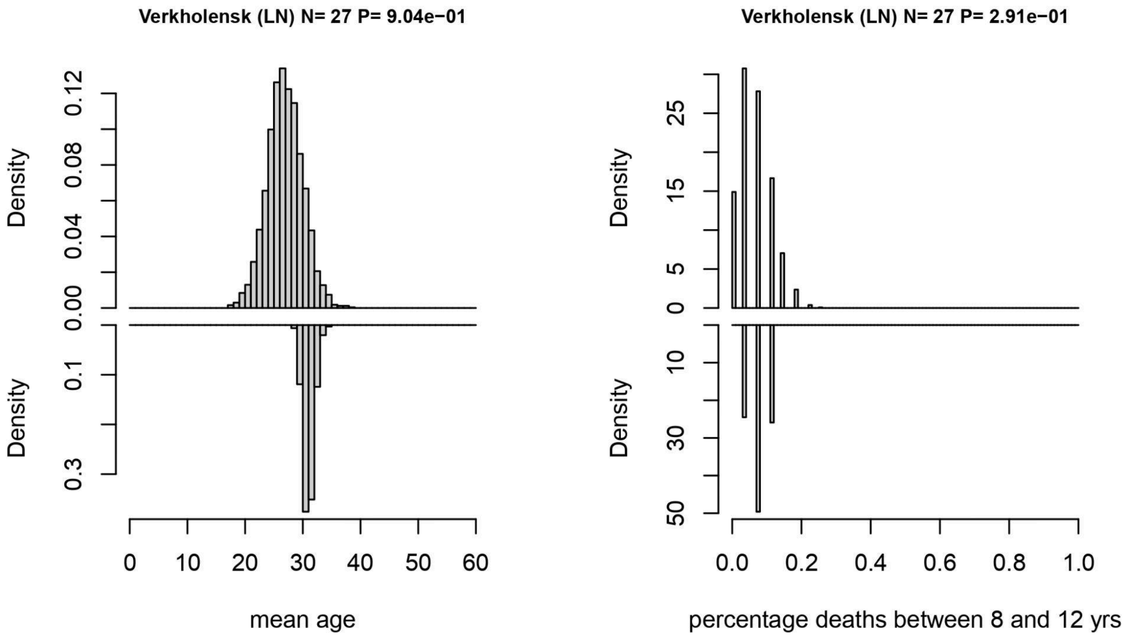


**Figure S22. Age-at-death modelling for Verkholensk.** Left: null distribution of mean age (top), test phase distribution of mean age (bottom). Right: null distribution of percentage of juvenile percentage (top), test phase distribution of juvenile percentage (bottom).
